# A real-time, transient kinetic study of *Drosophila melanogaster* Dicer-2 elucidates mechanism of termini-dependent cleavage of dsRNA

**DOI:** 10.1101/2020.09.29.319475

**Authors:** Raushan K. Singh, McKenzie Jonely, Evan Leslie, Nick A. Rejali, Rodrigo Noriega, Brenda L. Bass

## Abstract

*Drosophila melanogaster* Dicer-2 (dmDcr-2) differentially processes dsRNA with blunt or 2 nucleotide 3’-overhanging termini. We investigated the transient kinetic mechanism of these reactions using a rapid reaction stopped-flow technique and time-resolved fluorescence spectroscopy. We found that ATP binding to dmDcr-2’s helicase domain impacts the kinetics of dsRNA binding and dissociation in a termini-dependent manner, emphasizing the termini-dependent discrimination of dsRNA on a biologically-relevant time-scale. ATP-hydrolysis mediates local unwinding of dsRNA, and directional translocation on unwound single-stranded RNA, which is concurrent with a slow rewinding prior to dsRNA cleavage. Time-resolved fluorescence anisotropy reveals a nucleotide-dependent change in conformational dynamics of the helicase and Platform•PAZ domains in the nanosecond timescale that is correlated with termini-dependent dsRNA cleavage. Our study delineates kinetic events and transient intermediates for a Dicer-catalyzed reaction, thus establishing a framework for understanding other Dicers and how accessory factors modulate the reaction.

## INTRODUCTION

RNA silencing pathways play key roles in gene expression, heterochromatin formation, antiviral defense, and silencing of mobile genetic elements (Bartel 2018; Holoch and Moazed 2015; Obbard et al., 2009). In most cases these pathways involve an RNase III endonuclease called Dicer, which processes double-stranded RNA (dsRNA) precursors to produce small interfering RNAs (siRNAs) and microRNAs (miRNAs). These small dsRNAs have strands of ∼20-25 nucleotides (nts) and are subsequently passed to the RNA-Induced Silencing Complex (RISC), where they bind distinct Argonaute proteins to mediate silencing of target transcripts (Wilson et al., 2013).

*Caenorhabditis elegans* and *Homo sapiens* encode a single Dicer that produces both miRNAs and siRNAs. However, *Drosophila melanogaster*, and most arthropods (Jia et al., 2017), encode two Dicers. Dicer-1 is dedicated to the production of miRNAs, and Dicer-2 (dmDcr-2) produces siRNAs that silence endogenous as well as viral transcripts (Lee et al., 2004; Mussabekova et al., 2017). Like all metazoan Dicers, dmDcr-2 is a multidomain enzyme that includes an N-terminal helicase domain, a PAZ (Piwi•Argonaute•Zwille) domain, tandem RNase III domains, and a C-terminal dsRNA binding motif (dsRBM) (Figure S1). The enzyme is L-shaped with the helicase and PAZ domains at the base and top, respectively, and the RNase III domains at the center. The PAZ domain of dmDcr-2 shares structural homology with Argonautes, some of which show higher affinity for 2 nt 3’-overhanging (3’ovr) dsRNA termini compared to blunt (BLT) termini (Ma et al. 2004). As initially proposed (MacRae et al., 2006; Zhang et al., 2004), by interacting with the 3’ovr terminus, the PAZ domain allows Dicer to “measure” the distance to the RNase III domains, thus explaining the characteristic lengths of ∼20-25 base-pairs for siRNAs and miRNAs. For many years the PAZ domain and the adjacent Platform domain were considered the only Dicer domains that bound dsRNA termini (Zhang et al., 2004; MacRae et al., 2006; Park et al., 2011; Tian et al., 2014), but a recent cryo-EM structure of dmDcr-2 showed that in the presence of ATP, BLT dsRNA binds to the helicase domain (Sinha et al., 2018).

Innate immunity pathways have diverged dramatically during evolution, with antiviral defense involving an interferon response in vertebrates, and an RNA interference (RNAi) response in invertebrates (Schuster et al., 2019). Yet, pathways in both vertebrates and invertebrates involve Superfamily 2 (SF2) helicases of the same RIG-I-like receptor (RLR) subfamily (Fairman-William et al., 2010; Luo et al., 2013; Ahmad and Hur 2015) that in some cases show intriguing similarities. For example, molecular features at the termini of dsRNA are often used to distinguish viral and endogenous dsRNA, and the helicase domain of dmDcr-2 and the mammalian RLR, RIG-I, both have a higher binding affinity for dsRNA with BLT termini, a mimic of viral dsRNA termini, compared to 2 nt 3’ovr termini. Correspondingly, the ATPase activity for both proteins is activated to a greater extent by BLT, compared to 3’ovr, termini (Sinha et al., 2015; Marques et al., 2006; Ramanathan et al., 2016). Further, structural and biochemical studies show that the helicase domains of RIG-I and dmDcr-2 both change from an open to a more closed, or rigid, conformation when bound to dsRNA in the presence of nucleotide (Sinha et al., 2015; Kolakofsky et al., 2012).

While the helicase domains of dmDcr-2 and RIG-I are similar, the other domains of each protein have evolved in accordance with their roles in antiviral defense. Thus, although both helicase domains show similar activities, the more interesting question is how each helicase domain coordinates the activities of other domains. Biochemical and cryo-EM studies have provided evidence of diverse activities for dmDcr-2, including dsRNA binding, unwinding, translocation and cleavage, but how these reactions are coordinated and coupled to ATP hydrolysis is still unclear.

Understanding the mechanism of an enzyme-catalyzed reaction requires defining transient intermediates that are generated during the catalytic cycle, as well as the kinetic rates of interconversion between such states (Gutfreund 1995; Fersht 1999). Transient kinetic experiments are well-suited for delineating mechanistic details on a biologically relevant timescale (sec-min) (Williams 1991; Gutfreund 1995; Jencks 1997; Scholes et al., 2017). Towards these goals, we investigated the transient kinetic mechanism of the dmDcr-2-catalyzed reaction in real time using a stopped-flow system and time-resolved fluorescence spectroscopy. We find that ATP-binding to dmDcr-2 promotes initial recognition of dsRNA termini, and mediates clamping by the helicase domain. ATP hydrolysis induces local unwinding, followed by reannealing of dsRNA prior to cleavage by the RNase III domains; ATP hydrolysis fuels directional translocation of dmDcr2 on dsRNA until it arrives at the cleavage site. Importantly, our studies suggest an ATP-dependent modulation of the conformational dynamics of the helicase and Platform•PAZ domains, that in conjunction with the termini-dependent interactions with these domains, determines rates of dsRNA cleavage and siRNA release.

## RESULTS

### Kinetics of binding of dsRNA to dmDcr-2 is termini-dependent

Equilibrium binding experiments show that the affinity of dmDcr-2 for dsRNA depends on molecular features at the dsRNA terminus (Sinha et. al 2015). To identify transient intermediates of the enzyme•dsRNA interaction, and to gain insight into the kinetic basis of termini-dependent discrimination of dsRNA by dmDcr-2, we investigated the transient kinetic mechanism of dmDcr-2 binding to dsRNA using stopped-flow fluorescence experiments.

Cy3-end-labeled 52 base-pair dsRNAs (52-dsRNAs) with a BLT or 2 nt 3’ovr terminus were prepared by annealing top (sense) and bottom (antisense) strands as illustrated (Figures 1A and 1B; Methods). dmDcr-2 binds dsRNA at its termini, and one end of the dsRNA was blocked with two deoxynucleotides and biotin, to allow analysis only from the Cy3-labeled end. To simplify determination of kinetic parameters, stopped-flow experiments were performed under pseudo-first order conditions using 10-fold excess of dmDcr-2 over dsRNA. In all transient kinetic stopped-flow experiments with wild-type dmDcr-2 and mutant variants, initial binding events were kinetically well-separated from the dsRNA cleavage step, allowing kinetic analyses of binding without the need for blocking dsRNA cleavage.

**Figure 1:**
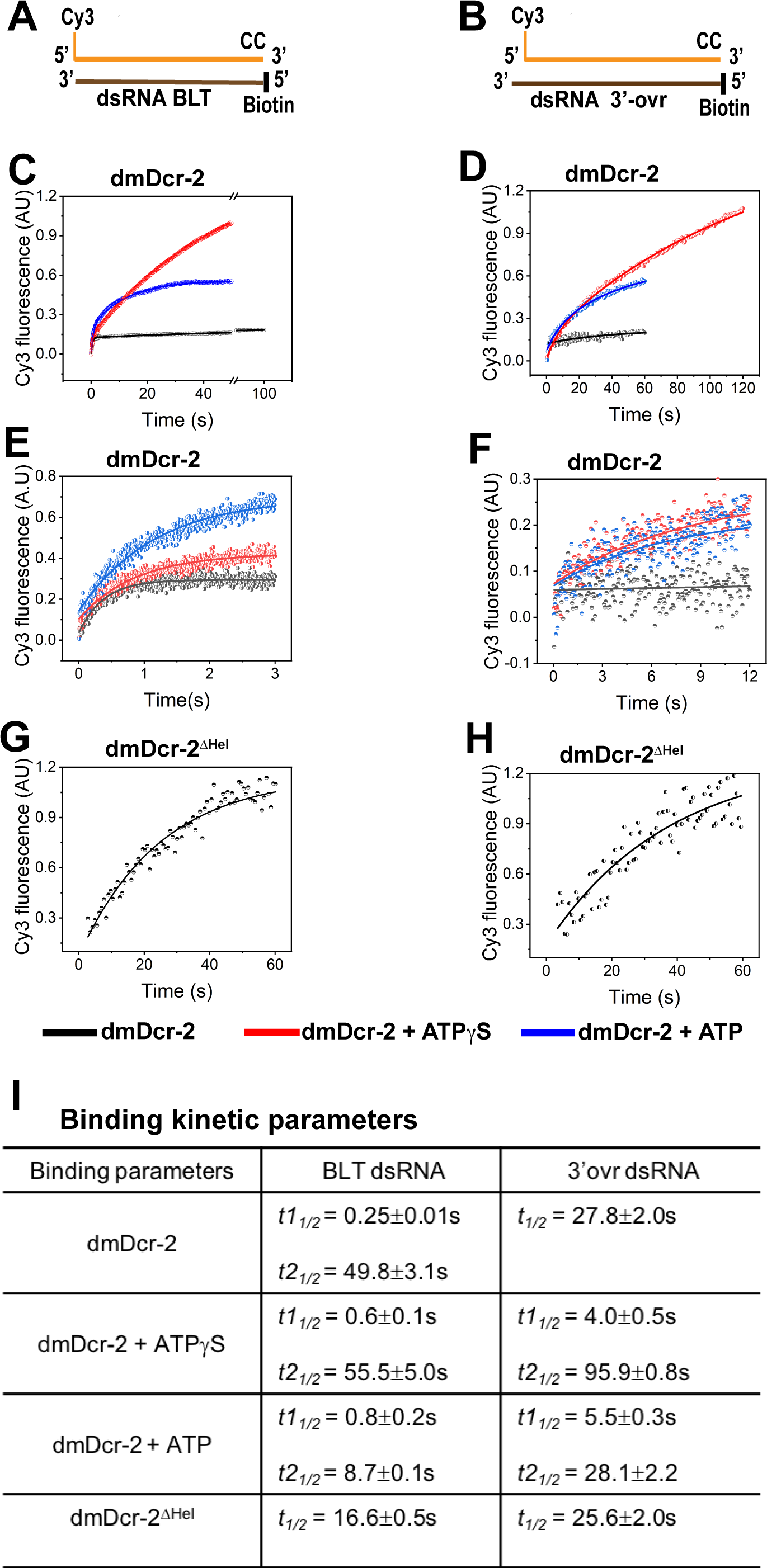
Stopped-flow kinetics of binding BLT and 3’ovr dsRNA to dmDcr-2. The time-dependent increase in the fluorescence signal of Cy3-end labeled 52-dsRNA (0.2µM) was monitored upon mixing with 10-fold excess of dmDcr-2 (2µM), alone and bound with nucleotide (ATPgS or ATP), in stopped-flow syringes. Cartoons show Cy3-end labeled 52-dsRNA with BLT **(A)** and 2 nt 3’ovr termini **(B)** with deoxynucleotides (CC) and biotin to prevent dmDcr-2 binding to one end. Representative kinetic traces for binding of Cy3-labeled BLT **(C)** and 3’ovr **(D)** 52-dsRNA to dmDcr2 are shown, with independent experiments over shorter time courses for BLT **(E)** and 3’ovr dsRNA **(F)**, as well as for binding of dmDcr-2^DHel^ to BLT (**G**) and 3’ovr (**H**) dsRNA. At least 4-5 traces were collected for each experimental condition, and averaged trace was analyzed with single or double exponential rate equations, yielding values for kinetic parameters (*kobs* = 0.693/t1/2) and associated standard error (**I**).

Equilibrium experiments indicate dmDcr-2 binds 3’ovr termini in the absence of nucleotide, while interactions with BLT termini requires ATP (Sinha et al., 2015). However, transient kinetic assays revealed interactions of both termini in the absence of nucleotide (black traces), and this was monitored on a long (Figures 1C and 1D) and short (Figures 1E and F) time scale. The BLT•dmDcr-2 complex may be less stable in the absence of nucleotide, possibly explaining why it was undetectable in prior equilibrium assays (Sinha et al., 2015). The kinetics of dmDcr-2 binding to BLT dsRNA were biphasic (*t11/2* =0.25s, *t21/2* =49.8s), but monophasic for binding to 3’ovr dsRNA (*t1/2* =27.8s) (Figure 1I). Prior studies suggest dmDcr-2 has two binding sites for dsRNA termini, one in the helicase domain and one in the Platform•PAZ domain (Sinha et al., 2015; Sinha et al., 2018). While previously reported equilibrium experiments indicate binding to the helicase domain is ATP dependent, we nevertheless considered the possibility that even in the absence of nucleotide, the biphasic kinetics reflected interactions with two binding sites. Indeed, experiments performed with dmDcr-2 lacking the helicase domain (dmDcr-2^DHel^), designed to delete one of the two binding sites, showed monophasic binding kinetics for both BLT and 3’ovr dsRNA (Figures 1G and 1H). Further, the half-life of binding to 3’ovr dsRNA in the absence of nucleotide was markedly similar for dmDcr-2 and dmDcr-2^DHel^ (Figure 1I) consistent with the idea that 3’ovr dsRNA is binding to the Platform•PAZ domain in both of these enzymes.

We further investigated the termini-dependent interactions of dsRNA by measuring the fluorescence-lifetime of the Cy3-end labeled dsRNA bound to dmDcr-2 (Figure S2; Methods). To avoid cleavage of dsRNA during the longer time scale required for time-resolved measurements, we used dmDcr-2 with a point mutation in each RNase III domain (dmDcr-2^RIII^; Figure S1). The dmDcr-2^RIII^•Cy3-BLT dsRNA complex showed two lifetimes (1.21ns and 2.28ns), but dmDcr-2^RIII^•Cy3-3’ovr dsRNA only one (1.28ns) (Figures S2C, S2D and S2G). Thus, in the absence of nucleotide, BLT dsRNA exists in two distinct microenvironments while bound to dmDcr-2, supporting the idea that the biphasic kinetics observed for BLT dsRNA binding to dmDcr-2 without nucleotide (Figures 1C and 1I) represents interactions with two sites.

Moreover, the fluorescence amplitude associated with the lifetime (τ Figure S2G) suggests that a major fraction (∼88%) of the BLT dsRNA is bound to the helicase domain, likely because of its higher binding specificity to this domain as compared to Platform-PAZ (Sinha et al; 2015; Marques et al., 2006; Jiang et al. 2011). Taken together, these data support the idea that in the absence of nucleotide, 3’ovr dsRNA binds to the Platform-PAZ domain of dmDcr-2 while BLT dsRNA binds poorly to this domain, and primarily to the helicase domain.

In the transient kinetic stopped-flow experiments (Figures 1C and 1D), addition of the non-hydrolyzable analog ATPgS (red trace) or ATP (blue trace), showed biphasic kinetics for binding of both BLT and 3’ovr 52-dsRNA to dmDcr-2 (Figure 1I), and this was more precisely measured over shorter times (Figures 1E and 1F). The fast-phase of binding was similar in the presence of either ATPgS or ATP, suggesting this phase is associated with ATP binding and unaffected by hydrolysis (Figure 1I). Further, the fast phase for binding BLT dsRNA (t1/2 = 0.6-0.8s) was an order of magnitude faster than binding to 3’ovr dsRNA (4.0-5.5s). In contrast to the initial fast phase of binding, the slow phase of binding kinetics became 6- and 3-fold faster, respectively, for BLT and 3’ovr dsRNA due to ATP hydrolysis (Figure 1I). Importantly, the half-life of this slow phase includes subsequent ATP hydrolysis-dependent steps, including unwinding/rewinding and translocation (see below).

While the biphasic binding kinetics observed in the absence of nucleotide appeared to derive from interactions with two binding sites, ATP binding has been associated with a conformational change in dmDcr-2’s helicase domain (Sinha et al., 2015; Sinha et al., 2018). Thus, we considered the possibility that in the presence of nucleotide the biphasic kinetics reflected an initial binding of dsRNA termini to the helicase domain, followed by a slow isomerization of the dmDcr-2•ATPgS•dsRNA complex. We tested this possibility by measuring the fluorescence lifetime of Cy3-end-labeled 52-dsRNA bound to dmDcr2^RIII^•ATPgS under equilibrium conditions (Figure S2). Only one fluorescence lifetime was observed, 2.14 and 1.31ns, respectively for BLT and 3’ovr (Figures S2E, S2F and S2G) –ruling out the possibility of two binding sites in the presence of ATPgS, and indicating the presence of a slow isomerization step upon initial binding of dsRNA termini to dmDcr-2’s helicase domain. These data indicate that in the absence of nucleotide, dsRNA interacts with two binding sites in dmDcr-2, in a termini-dependent manner. However, in the presence of ATPgS, BLT and 3’ovr dsRNA predominantly bind to a single binding site (helicase domain), albeit with a 6- and 2-fold difference in half-lives, respectively, for the fast and slow phases of binding (Fig 1I). Thus, the two termini show distinct macromolecular interactions within the helicase domain.

### Blunt dsRNA has a higher residence time in the dmDcr-2•ATPgS complex than 3’ovr dsRNA

Our prior studies show that dmDcr-2 cannot hydrolyze ATPgS (Sinha et al., 2015), and the use of this analog allowed us to trap an important reaction intermediate: dsRNA interacting with the helicase domain (Sinha et al., 2018). Taken together, the studies described above (Figures 1 and S2) indicate the initial encounter complex (dmDcr-2•ATPgS•dsRNA) slowly isomerizes to a final stable complex (dmDcr-2*•ATPgS•dsRNA). Further, the half-life values of these transient steps (Figure 1I, ATPgS) differ for BLT and 3’ovr dsRNA indicating that the initial encounter and the isomerization are termini dependent. We anticipated that these termini-dependent interactions would impact the dissociation off-rates (residence time), which in turn could impact termini-dependent processivity of dmDcr-2 on long dsRNA. To gain insight into the residence time, and the stability and energetics of termini-dependent interactions of dsRNA with the helicase domain, we measured dissociation off-rates for dsRNA bound to dmDcr-2 in the presence of ATPgS using the stopped-flow system.

**Figure 2:**
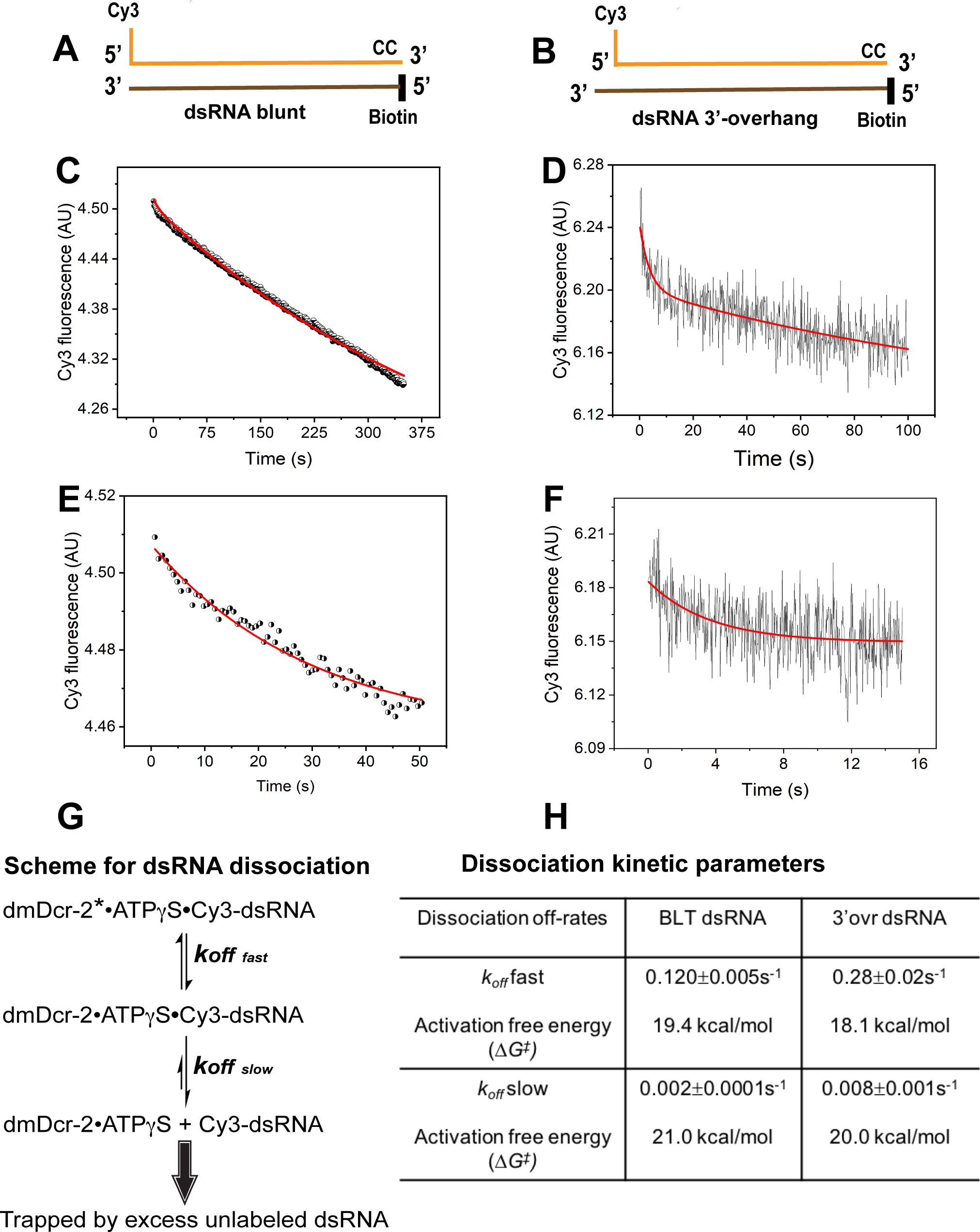
Stopped-flow kinetics for dissociation of BLT and 3’ovr dsRNA bound to dmDcr-2•ATPgS. Dissociation kinetics were measured by mixing enzyme•bound Cy3-end labeled 52-dsRNA with 10-fold excess of unlabeled dsRNA in stopped-flow syringes, and monitoring decrease in Cy3 fluorescence over time. Cartoons show BLT (A) and 3’ovr (B) dsRNA with representative kinetic traces for dissociation of Cy3-labeled BLT **(C)** and 3’ovr **(D)** 52-dsRNA, and independent experiments over shorter time courses (BLT, **E**; 3’ovr **F)**. (**G**) Reaction scheme for dissociation of enzyme-bound Cy3-dsRNA. **(H)** Kinetic traces were analyzed with single or double exponential rate equations, yielding values for dissociation off-rates. At least 4-5 traces were collected for each condition, and averaged trace was analyzed with single or double exponential rate equations to obtain values for kinetic parameters (*kobs* = 0.693/t1/2) and associated standard error.

We prepared the dmDcr-2*•ATPgS•Cy3-dsRNA complex by incubating Cy3-end-labeled dsRNA (Figures 2A and 2B) with saturating concentrations of dmDcr-2 and nucleotide for 5 minutes (to allow formation of stable complex), and then mixed with 10-fold excess of unlabeled dsRNA to trap dissociated dmDcr-2•ATPgS (Figure 2G). Dissociation of both BLT and 3’-ovr dsRNA in the presence of ATPgS was biphasic (Figures 2C and 2D), as shown more precisely over a shorter time scale (Figures 2E and 2F). The two-step dissociation of dsRNA (biphasic kinetics) is consistent with the two-step binding reaction of dsRNA with dmDcr-2•ATPgS (Figure 1I), as expected from the principle of microscopic reversibility (Krupka et al., 1966).

Analyses with a double exponential rate equation yielded values for *koff* (dissociation off-rate; Figure 2H). While values for fast and slow *koff* for BLT were 0.12s^-1^ and 0.002s^-1^ respectively, dissociation of 3’ovr was 2 and 4-fold faster, with fast and slow koff values of 0.28s and 0.008s -1 -1 (Figure 2H). This confirmed that BLT dsRNA was engaged more tightly than 3’ovr dsRNA in the dmDcr2•ATPgS complex. Activation free energy barriers (DG^‡^), calculated using the Eyring equation (see Methods), for dissociation of BLT dsRNA were 1.3 and 1.0 kcal/mol higher, respectively, for fast and slow phase dissociation compared to values for 3’ovr dsRNA (Figure 2H). This emphasizes that the molecular forces involved in holding BLT termini in the helicase domain of dmDcr-2 bound with ATPgS are stronger than those for 3’ovr termini. Notably, a slower dissociation off-rate of the enzyme-bound BLT dsRNA, in comparison to the observed rate of its cleavage (see below), is consistent with the observed processivity with this substrate (Welker et al., 2011; Sinha et al., 2015; Sinha et al., 2018). In other words, dmDcr-2 has a higher probability of cleaving BLT dsRNA (as compared to 3’ovr) before it dissociates from the productive enzyme-substrate complex.

### dmDcr-2 catalyzes an ATP-dependent unwinding and rewinding of dsRNA

dmDcr-2 belongs to the SF-2 family of helicases, many of which catalyze an ATP-dependent directional unwinding of duplex RNA or DNA (Fairman-Williams et al., 2010). The dmDcr-2 helicase domain is similar to that of mammalian RIG-I, and while there are conflicting reports (Takahasi et al., 2007; Myong et al., 2009), RIG-I has been reported to lack unwinding activity (Myong et al., 2009). Thus, the observation of a single-stranded RNA within the helicase domain in the cryo-EM structure of dmDcr-2•ATPgS•BLT-dsRNA is notable (Sinha et al., 2018). Biochemical assays based on strand-displacement confirmed unwinding activity (Sinha et al., 2018), but raised the question of why dmDcr-2 unwinds dsRNA, since presumably it would need to rewind the dsRNA for subsequent cleavage by the dsRNA-specific RNase III active sites (Nicholson 2014).

To directly monitor ATP-dependent unwinding of dsRNA catalyzed by dmDcr-2 in real time, and to test whether rewinding occurs, we performed stopped-flow experiments utilizing BLT and 3’ovr 52-dsRNA that contained a Cy3-Cy5 donor–acceptor Förster resonance energy transfer (FRET) pair at one terminus (Figures 3A and 3B). We anticipated that unwinding and rewinding of Cy3-Cy5 labeled dsRNA by dmDcr-2 would produce a sequential loss and gain in FRET signal over time. While dmDcr-2 alone (black) did not produce a detectable change in the FRET signal of BLT or 3’ovr dsRNA, a biphasic change in FRET as a function of time was observed in the presence of ATP. The half-lives for ATP-dependent unwinding of BLT and 3’ovr were 0.5s and 1.0s, respectively (Figure 3E), which correlates with the 2-fold difference in the rate of ATP hydrolysis by dmDcr-2 while bound to these dsRNAs (Sinha et al., 2015). Interestingly, the kinetic parameters for dmDcr-2-catalyzed unwinding are much slower than those of processive helicases involved in replication (Bianco et al., 2001; Burnham et al., 2019), and more similar to the ATP-dependent unwinding by non-processive helicases, such as, *E. coli* Rep helicase (Bjornson et al., 1994; Ha et al., 2002).

**Figure 3:**
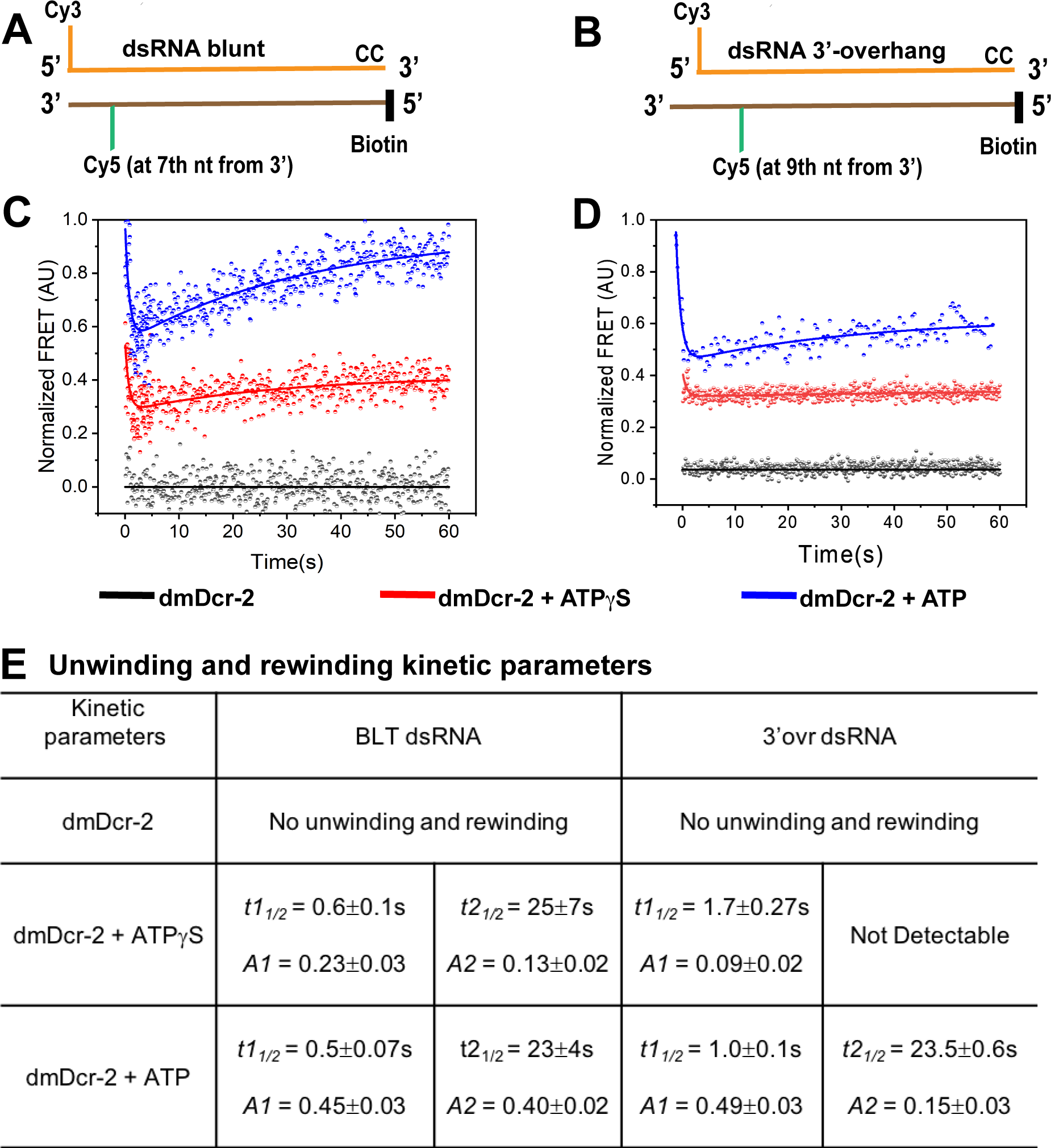
Stopped-flow kinetics of dsRNA unwinding and reannealing by dmDcr-2. The time-dependent change in FRET signal was monitored after mixing 0.2µM dsRNA containing a FRET (donor-acceptor) pair with 2µM dmDcr-2 in stopped-flow syringes. Cartoons showing BLT (**A**) and 3’ovr (**B**) 52-dsRNA indicating positions of Cy3 and Cy5, and modifications to block dmDcr-2 entry, as in Figure 1. Representative kinetic traces for unwinding and reannealing of FRET-labeled BLT (**C**) and 3’ovr (**D**) dsRNA by dmDcr-2 alone (black), with ATPgS (red) or ATP (blue). Kinetic traces were analyzed with a double exponential rate equation, and kinetic parameters are in (**E**). At least 4-5 traces were collected for each experimental condition, and the averaged trace was analyzed with single or double exponential rate equations, yielding values for kinetic parameters (*kobs* = 0.693/*t1/*2), amplitude (A), and associated standard error.

To understand the properties of kinetic intermediates of the dsRNA unwinding/rewinding reaction, we compared the amplitudes associated with distinct phases of the reaction (Figure 3E). The amplitudes of the unwinding (*A1*) and rewinding (*A2*) phases in the presence of ATP were almost the same for BLT dsRNA, that is, the FRET associated with the unwound state returned to the value observed in the initial annealed state. By contrast, for the 3’ovr dsRNA, the amplitude for the rewinding phase was only 30% of the unwinding value. This feature suggested that unlike BLT dsRNA, the 3’ovr dsRNA remained partially unwound at the end of this assay. As observed for human Dicer, possibly the unwound state is stabilized by interactions with the Platform•PAZ domain prior to cleavage (Park et al., 2011; Tian et al., 2014). In this scenario BLT would interact weakly with the Platform•PAZ domain, without unwinding of its terminal base-pairs.

We also observed a sequential loss and gain of FRET signal catalyzed by dmDcr-2 on BLT and 3’ovr with ATPgS, although the signal was not as robust as with ATP. This result may indicate the presence of contaminating ATP in the commercially-purchased ATPgS, although we cannot rule out the possibility that binding of nucleotide to dmDcr-2 (without hydrolysis) is sufficient to initiate unwinding, as reported for some DEAD-box helicases (Liu et al., 2008). However, with BLT dsRNA we observed both unwinding and rewinding, which is more consistent with the presence of contaminating ATP; since ATP hydrolysis is ∼2 fold less efficient with 3’ovr dsRNA compared to BLT dsRNA (Sinha et al. 2015), as might be expected, rewinding was detectable in the presence of ATPgS for BLT, but not for 3’ovr dsRNA. With the addition of ATP, rewinding was detectable for 3’ovr dsRNA, and we observed an almost 3-fold higher amplitude (*A2*) associated with rewinding for BLT dsRNA compared to ATPgS, underscoring the critical role of ATP-hydrolysis in the rewinding reaction (Yusufzai et al., 2008; Yusufzai et al., 2010).

### ATP-dependent translocation of dmDcr-2 on dsRNA poises the enzyme for RNA cleavage

The ATP-dependent unwinding activity of helicases is often coupled with their processive translocation on single or double-stranded nucleic acid (Singleton et al., 2007; Ha et al., 2012). Indeed, our prior end-point assays using BLT dsRNA with steric blocks indicate dmDcr-2 translocates dsRNA through the helicase domain (Sinha et al., 2018). We sought to acquire direct, real-time transient kinetic information for ATP-dependent translocation of dmDcr-2 to the cleavage site, and to correlate the speed of translocation with termini-dependent differences in dsRNA cleavage rates.

The major dmDcr-2 cleavage products are siRNAs with strands of 21 or 22 nts (Zamore et al, 2000; Elbashir et al., 2001). To study the ATP-dependent translocation of dmDcr-2 to the predominant cleavage site in real time, we monitored the protein-induced fluorescence enhancement (PIFE) signal of Cy3 attached to the 18^th^ position of the sense strand of 52-dsRNA (Figure 4A and 4B). Representative stopped-flow kinetic traces showed the time-dependent increase in Cy3 fluorescence signal in the presence of ATP (Figures 4C and 4D), and analysis with a single-exponential rate equation yielded half-life values for the 18-base-pair translocation of dmDcr-2 on BLT and 3’ovr dsRNA, of 14s and 27.8s, respectively (Figure 4E). The 2-fold higher translocation rate of dmDcr-2 on BLT dsRNA compared to 3’ovr dsRNA is consistent with the 2-fold higher rate of ATP hydrolysis observed in the presence of BLT dsRNA compared to 3’ovr dsRNA (Sinha at al., 2015), supporting the idea that the free-energy of ATP hydrolysis is directly coupled to translocation. These values are similar to the average duration (18.5s) of ATP-dependent translocation catalyzed by RIG-I on a 25-base-pair BLT dsRNA (Myong et al., 2009), suggesting the kinetic mechanism of ATP-dependent translocation by these two helicases is similar.

**Figure 4:**
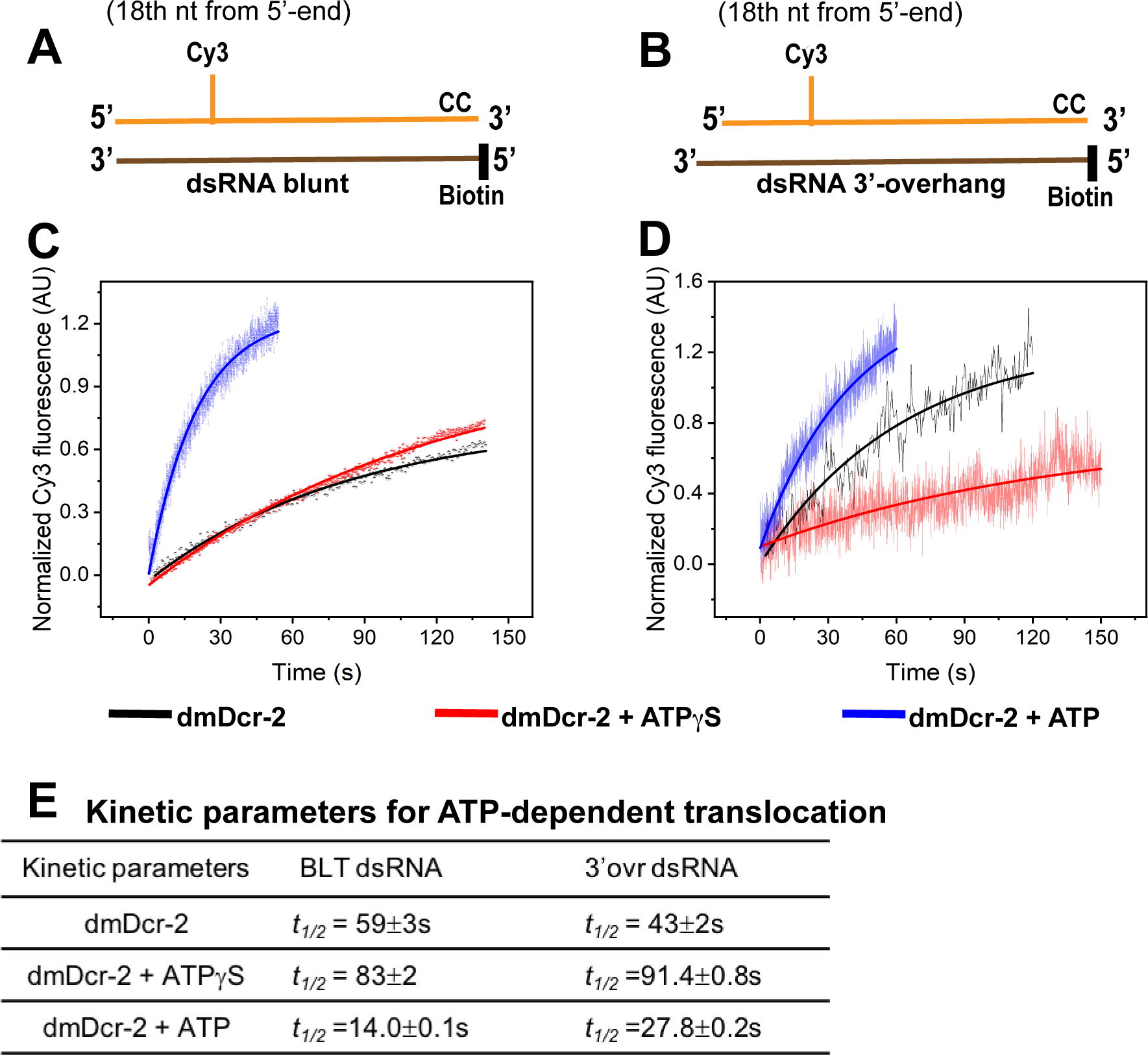
Stopped-flow kinetics for ATP-dependent translocation of dmDcr-2 on BLT and 3’ovr dsRNA. Cartoons show BLT (**A**) and 3’ovr (**B**) 52-dsRNA with a covalently-linked Cy3 at the 18^th^ nucleotide of the top (sense) strand, and deoxynucleotide (CC) and biotin to prevent binding of dmDcr-2 to one end. Representative stopped-flow kinetic traces for translocation of dmDcr-2 alone (black trace), and while bound with ATPgS (red) and ATP (blue) on BLT (**C**) and 3’ovr (**D**) dsRNA. Kinetic traces were analyzed with a single exponential rate equation, yielding observed rate constants associated with translocation (**E**). At least 4-5 traces were collected for each experimental condition, and averaged trace was analyzed with single or double exponential rate equations, yielding kinetic parameters (*kobs* = 0.693/*t1/*2) and associated standard error.

We also observed a time-dependent increase in Cy3 fluorescence in our translocation assay with ATPgS (red trace) or even without nucleotide (black trace) (Figure 4C and 4D), albeit the rates and associated signal amplitudes were markedly lower compared to those in the presence of ATP (Figure 4E). Consistent with our binding studies (Figures 1 and S2), in the absence of ATP, 3’ovr dsRNA does not thread through the helicase domain, but directly interacts with the Platform•PAZ domain, which presumably directs the dsRNA to the RNase III active sites for cleavage. Therefore, it seems likely that the enhancement in fluorescence from Cy3 at position 18 of 3’ovr dsRNA in the absence of nucleotide reflects an interaction with the protein near the RNase III domains. Similarly, in the presence of ATPgS, the observed signal enhancement may reflect that the stable intermediate observed with this analog is proximal to the Cy3 at position 18. Taken together, these kinetic studies demonstrate that dmDcr-2 utilizes the free energy of ATP hydrolysis to translocate along the sense (top) strand from 5’ to 3’, as observed for RIG-I (Myong et al., 2009; Devarkar 2018).

### ATP-dependent cleavage of dsRNA is dictated by dsRNA termini

After delineating the transient kinetic events of the dmDcr-2-catalyzed reaction preceding the ATP-dependent cleavage (Figures 1-4), we wished to gain mechanistic information about how these kinetic events fine-tune the culmination of the reaction, dsRNA cleavage and siRNA release. Towards this goal, we performed transient kinetic experiments to directly monitor substrate cleavage and product release in real time. Notably, we found that the dmDcr-2-catalyzed dsRNA cleavage/siRNA release was kinetically well separated from earlier kinetic events (binding, unwinding/rewinding and translocation), enabling reliable evaluation of these parameters (see below and Figures 1, 3 and 4).

We prepared BLT and 3’ovr 52-dsRNA with a Cy3-Cy5 FRET pair that spanned the primary cleavage site in 52-dsRNA (Figures 5A and 5B, dotted lines), so that dmDcr-2-catalyzed cleavage coupled with siRNA release would lead to a loss of FRET. We mixed 10-fold excess of dmDcr-2 (alone or with ATP or ATPgS) with the FRET-labeled 52-dsRNA in the stopped-flow system and monitored the reaction for ∼5 min to observe the entire catalytic cycle. We recorded real-time progress of the reaction by monitoring the time-dependent change in Cy5 FRET signal at 670nm upon excitation of the donor Cy3 at 530nm.

**Figure 5:**
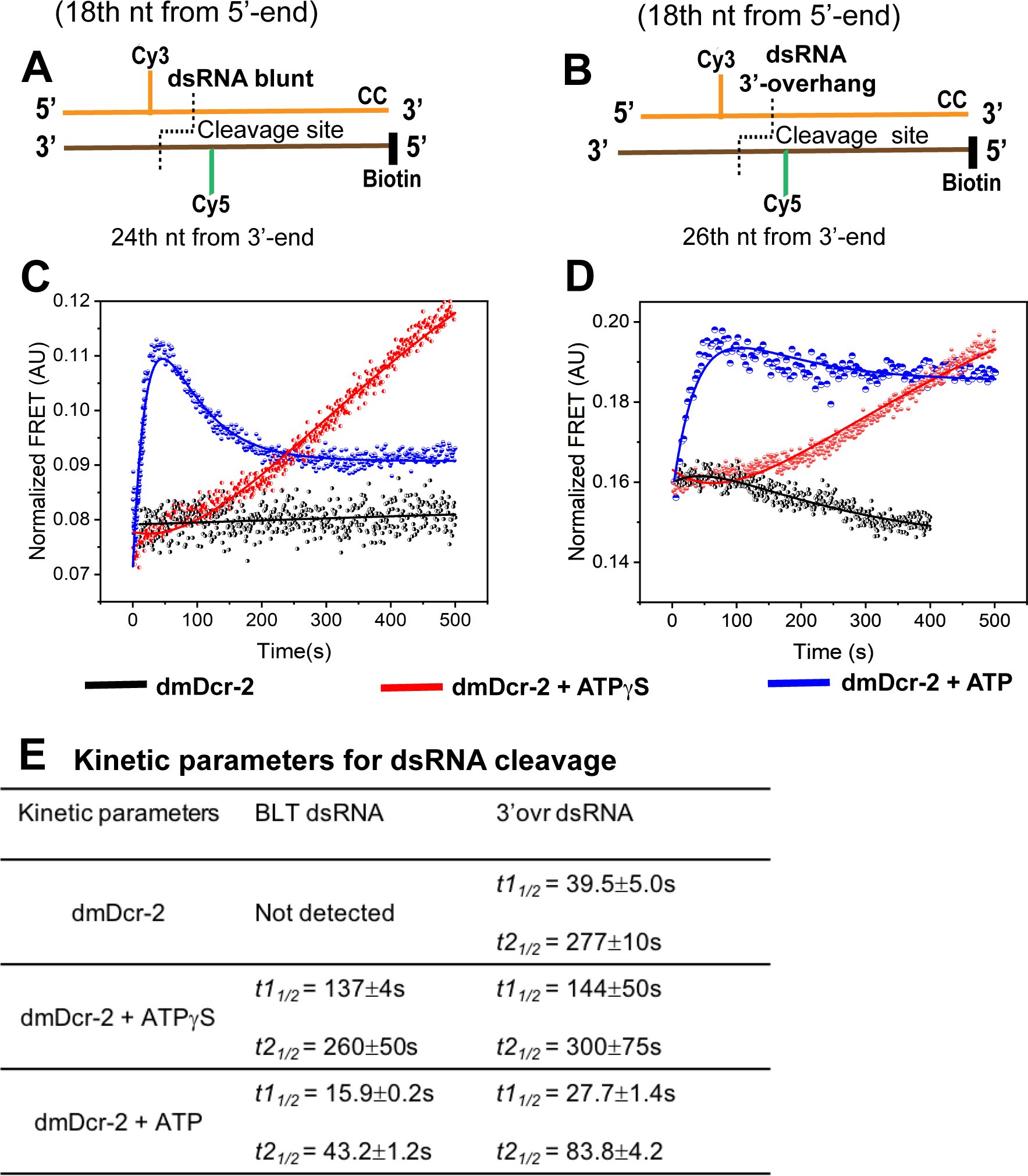
Stopped-flow kinetics for the cleavage of BLT and 3’ovr 52-dsRNA by dmDcr-2. Cartoons illustrate BLT (**A**) and 3’ovr (**B**) dsRNA containing the Cy3-Cy5 FRET pair at positions indicated, and deoxynucleotide (CC) and biotin to block dmDcr-2 binding at one end. Representative stopped-flow kinetic traces for cleavage of BLT (**C**) and 3’ovr (**D**) 52-dsRNA by dmDcr-2 alone (black), with ATPgS (red) or ATP (blue). Kinetic traces were analyzed with a double exponential rate equation, and kinetic parameters listed in (**E**). At least 4-5 traces were collected for each condition, and averaged trace was analyzed with single or double exponential rate equations, yielding values for kinetic parameters (*kobs* = 0.693/t1/2) and associated standard error.

dmDcr-2 alone did not produce a significant change in FRET signal for BLT dsRNA, but biphasic kinetics were observed for 3’ovr dsRNA in the absence of nucleotide, with an initial modest gain of FRET signal followed by a slow loss over time (Figures 5C and 5D, black traces), as expected for substrate binding followed by slow cleavage. This result is consistent with prior studies indicating that dmDcr-2’s Platform•PAZ domain mediates an ATP-independent cleavage of 3’ovr dsRNA, but not BLT dsRNA (Sinha et al., 2015; Sinha et al., 2018). Interestingly, we observed a similar trace for cleavage of BLT dsRNA by dmDcr-2 lacking the helicase domain (dmDcr-2^DHel^) (Figure S3C, cyan trace), likely mediated via its binding to the Platform-PAZ domain in the absence of competitive binding to the helicase domain. Not surprisingly, the rate of 3’ovr cleavage catalyzed by dmDcr-2^DHel^ was ∼2.5 fold higher than BLT (Figure S3C, S3D and S3E)— supporting the idea that a lower binding specificity of BLT (as compared to 3’ovr) for the Platform-PAZ domain impacts the rate of dsRNA cleavage (Ma et al., 2004).

In the presence of a non-hydrolyzable analog of ATP (ATPgS), both BLT and 3’ovr dsRNA showed biphasic kinetics (Figures 5C and 5D, red trace) where both phases were associated with an increase in the FRET signal, suggesting that these dsRNAs are kinetically trapped in a non-productive conformation by the helicase domain in the absence of ATP hydrolysis. In contrast, in the presence of ATP, both BLT and 3’ovr showed biphasic kinetics with an initial fast increase of FRET, followed by a slow loss as a function of time.

In an analogous manner to the PIFE observed in the translocation assay (without FRET acceptor, Figures 4C and 4D), we anticipated that an increase in Cy3 fluorescence due to PIFE, would also lead to increased Cy5 fluorescence due to a greater extent of energy transfer from Cy3 to Cy5 (Stennett et al., 2015; Nguyen et al., 2019). Indeed, the time-dependent increase in FRET signal observed for BLT and 3’-ovr dsRNA in the presence of dmDcr-2 and ATP was similar to the increase in PIFE signal observed in the translocation assay (compare t1/2 values with ATP in Figure 4E and 5E).

Upon completion of the fast phase (FRET increase) for reactions containing dmDcr-2 and ATP, we observed a slow loss of FRET over time with both BLT and 3’ovr dsRNA (blue traces, Figures 5C and 5D), which we attributed to cleavage of dsRNA coupled with siRNA release. To confirm this interpretation, we performed the FRET-based assay with dmDcr2^RIII^ to preclude cleavage (Figure S4). Only the time-dependent increase in FRET was observed, confirming that the loss of FRET signal in the presence of ATP correlates with dsRNA cleavage and siRNA release. The half-life for the ATP-dependent cleavage/product release for BLT dsRNA (*t1/2* = 43s) was two-fold faster than that of 3’-ovr dsRNA (*t1/2* = 83s) (Figure 5E). The termini-dependent difference in cleavage/release rates in the presence of ATP could be due to a termini-dependent interaction of dsRNA after translocation to the Platform•PAZ domain, which poises the enzyme for substrate cleavage (see below). The 2-fold difference in dsRNA cleavage and siRNA release also correlates with the difference in ATP-hydrolysis rate observed with dsRNA with different termini (Sinha et al., 2015). Further, a markedly higher amplitude for the FRET loss during BLT cleavage/siRNA release (as compared to 3’ovr) indicates an incomplete release and/or rebinding of 3’ovr siRNA. This feature could arise due to a higher binding specificity of 3’ovr to Platform•PAZ domain, and thus play a critical role in siRNA release (see below) (Ma et al., 2004; Gan et al., 2006; Nicholson 2014).

### Termini-dependent conformational dynamics of helicase and PAZ•platform domains impact catalysis

Conformational dynamics/fluctuations of enzymes in the nanosecond time scale are often linked with their substrate specificity and catalytic turnover rate (Broos et al., 1995; Henzler-Wildman et al., 2007; Banerjee and Pal, 2008). When covalently attached to DNA or RNA, changes in the fluorescence properties of probes such as Cy3 and 2-Aminopurine can be used to investigate conformational dynamics of nucleic acid binding proteins (Luo et al., 2007; Dunlap et al., 2002; Purohit et al., 2003). Our transient kinetics studies indicated that, in the presence of ATP, BLT and 3’ovr termini are discriminated during initial binding, and prior to their cleavage, respectively, by the helicase and Platform•PAZ domains (Figures 1 and 5). To gain information about termini-dependent differences in conformational dynamics of these domains that could be correlated with the catalytic activity of the enzyme, we performed time-resolved fluorescence anisotropy using Cy3-labeled dsRNA. We measured anisotropy parameters of Cy3-end labeled BLT and 3’ovr 52-dsRNA (Figures 6A and 6B), alone (green), bound to dmDcr-2 (black), and bound to dmDcr-2 with ATPgS (red) or ATP (blue). Again, to avoid dsRNA cleavage during the longer times required for time-resolved measurements, we used dmDcr-2^RIII^, which binds and hydrolyzes ATP similar to the wildtype enzyme (Cenik et al., 2011). All incubations were ∼5 minutes so that kinetic events prior to dsRNA cleavage were complete (one catalytic turnover requires 1-2 minutes; Fig 5E). Anisotropy decay curves (Figure 6C-J) were analyzed with single and double exponential rate equations yielding values of rotational correlation time (φ), fractional amplitude (*A*), and limiting anisotropy (r*_∞_*) (see Methods and Figure 6K). The change in anisotropy (Dr) was evaluated by subtracting the value of limiting anisotropy (Δr) from the fundamental/initial anisotropy (r0) (Lakowicz 2006).

**Figure 6.**
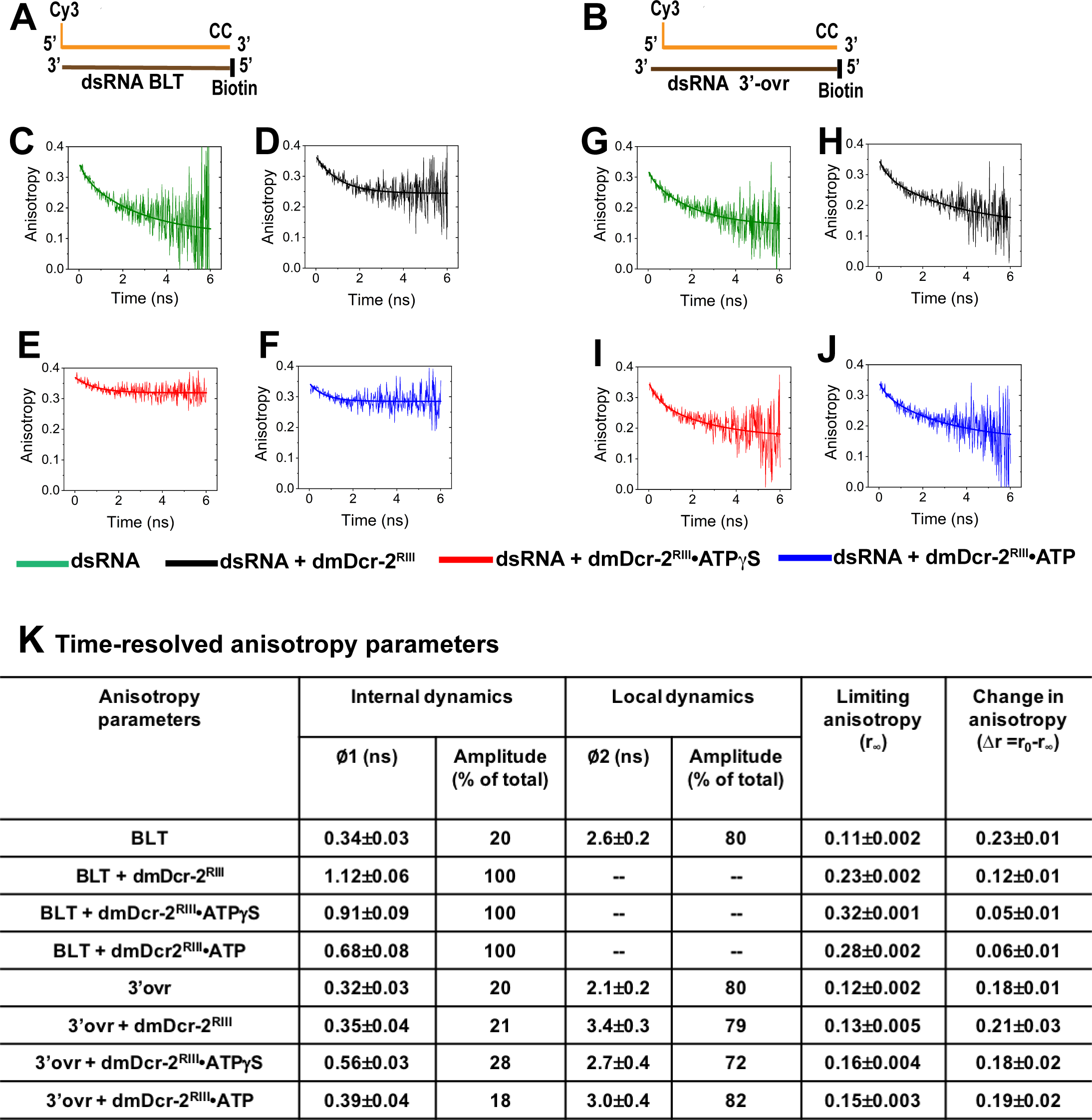
Time-resolved fluorescence anisotropy of dmDcr-2 bound to BLT (**A**) and 3’-ovr (**B**) 52-dsRNA reveals conformational dynamics of helicase and Platform•PAZ domains. Representative anisotropy decay curves for Cy3-end labeled BLT **(C-F)** and 3’ovr dsRNA **(G-J)** alone (green) and bound to dmDcr-2^RIII^ in the absence of nucleotide (black), or in the presence of ATPgS (red) or ATP (blue). All decay curves were analyzed with a minimum number of exponential terms (single and double exponential rate equations) to obtain the best fit (see Methods and Figure S5), yielding values of anisotropy parameters **(K)**. The precise values of anisotropy parameters associated with the fast correlation time (φ1) were determined from independent analyses of the fast phase of anisotropy decay curves using a single exponential equation (see Figure S5).

The anisotropy parameters for Cy3-end labeled BLT and 3’ovr 52-dsRNA (Figures 6C and 6G) were similar to each other and to reported values of Cy3 end-labeled dsDNA (Sanborn et al., 2011). To obtain precise values for the fast correlation times, the fast phase of anisotropy decay was analyzed separately (Figure S5A-J). Fast and slow rotational correlation times for BLT 52-dsRNA were 0.34 and 2.6ns, respectively, with associated amplitudes of 20% and 80% (Figures 6K). For Cy3-end labeled DNA and RNA, fast correlation times derive from internal rotational dynamics of the fluorophore, while slow correlation times report on local tumbling of the hydrated nucleic acid (Unruh et al., 2005). The similar anisotropy parameters of Cy3-end labeled BLT and 3’ovr 52-dsRNA (Figures 6C, 6G and 6K) indicate internal dynamics of the terminal Cy3, and local tumbling of the dsRNAs, were similar in our experimental conditions.

Binding kinetics and fluorescence lifetime measurements showed that BLT termini are primarily engaged with the helicase domain in the presence of ATPgS (Figures 1 and S2). In view of revealing conformational dynamics of the helicase domain while engaged with BLT termini, we compared anisotropy parameters of BLT 52-dsRNA alone (Figure 6C) to those obtained in the presence of dmDcr-2^RIII^ (Figure 6D) and dmDcr-2^RIII^•ATPgS (Figure 6E; Figure 6K). Notably, the binding of enzyme to BLT dsRNA in the absence of nucleotide diminished the second phase of the anisotropy decay curve, consistent with a highly restricted local tumbling of enzyme-bound BLT dsRNA. Further, the fast correlation time associated with internal rotational dynamics of Cy3 attached to BLT dsRNA became longer by ∼3.5-fold upon binding to dmDcr-2’s helicase domain (Figure 6K), suggesting a markedly reduced conformational fluctuation of the helicase domain upon engagement with BLT termini even without nucleotide. The above marked reduction in the fast correlation time (internal dynamics) of BLT dsRNA upon binding to dmDcr-2 is consistent with enhanced molecular rigidity in close proximity to the termini, leading to an increase in fluorescence life-time of Cy3 (Figure S2G). The presence of ATPgS had a modest effect on the fast correlation time when compared to protein alone, but the anisotropy change (Dr) was reduced by ∼2.4 fold, again consistent with a nucleotide-mediated rigidification of the helicase domain (Figures 6D, 6E and 6K). These data corroborate the rigid/closed conformation of dmDcr-2•ATPgS•BLT predicted by limited proteolysis and observed by cryo-EM (Sinha et al., 2015; Sinha et al., 2018).

Our binding kinetics and time-resolved fluorescence measurements indicated that, like BLT dsRNA, in the presence of ATPgS, the 3’ovr terminus is engaged with the helicase domain (Figure 1 and S2). To get insight into termini-dependent modulation of helicase dynamics, we compared anisotropy parameters of 3’ovr and BLT dsRNA in the dmDcr-2•ATPgS complex. In contrast to experiments with BLT dsRNA, we observed both internal dynamics, as well as local tumbling of 3’ovr (Figures 6G-K). The internal dynamics, when comparing 3’ovr dsRNA alone (0.32ns) with values for addition of dmDcr-2^III^ and ATPgS (0.56ns), became ∼1.75 fold slower (Figures 6K and S5). This feature is consistent with a small change in molecular rigidity at the 3’ovr terminus as compared to BLT upon binding to dmDcr-2, manifested as a modest change in the fluorescence life-time (Figure S2G). Further, the change in anisotropy associated with BLT bound to dmDcr-2•ATPgS (Dr, 0.05) is ∼3.5 fold lower than that of 3’ovr (Dr, 0.18; Figure 6K), consistent with the ∼4-fold slower dissociation off-rate of enzyme-bound BLT dsRNA as compared to 3’ovr dsRNA in the presence of ATPgS (Figure 2).

Prior studies (Cenik et al., 2011; Sinha et al 2015), as well as observations reported here (Figure 5), indicate ATP hydrolysis is required for cleavage after dsRNA engages the helicase domain, and the non-hydrolyzable analog ATPgS traps both BLT and 3’ovr termini within dmDcr-2’s helicase domain. With the addition of ATP, hydrolysis fuels translocation and enables interaction of the dsRNA termini with the Platform•PAZ domain, to promote “measuring” and cleavage of the dsRNA to produce 21-22 nt siRNAs (Figures 4 and 5). To specifically compare interactions of BLT and 3’ovr termini with the Platform•PAZ domain, we compared anisotropy parameters for BLT and 3’ovr dsRNA in the presence of dmDcr-2^RIII^ and ATP. As for BLT termini bound to the helicase domain (i.e. with ATPgS; Figure 6E), the long correlation time was absent for BLT dsRNA interacting with the Platform•PAZ domain (i.e. with ATP; Figures 6F and 6K). In contrast, both fast and slow correlation times were present for 3’ovr in the presence of ATP (Figure 6J), and further, the fast correlation time was shorter than observed for BLT termini (f1, 0.39ns, 3’ovr; 0.68ns, BLT, Figure 6K), indicating the conformational fluctuation of the Platform•PAZ domain in a catalytically competent state is ∼1.75-fold lower when bound to BLT, compared to 3’ovr, termini. The higher local rigidity of the Platform•PAZ domain at the BLT terminus in the presence of ATP (as compared to 3’ovr) is also consistent with a markedly higher increase in fluorescence life-time of Cy3-BLT dsRNA upon binding to dmDcr-2^RIII^ •ATP, which is caused by an increase in molecular rigidity near the fluorophore, as compared to that of 3’ovr (Figure S6). Since the ATP-dependent processing of BLT dsRNA (cleavage and siRNA release) is ∼2-fold faster than 3’ovr (Figure 5E), the reduced conformational fluctuation of the Platform•PAZ domain is correlated with enzyme catalysis (Broos et al., 1995; Henzler-Wildman and Kern., 2007). Interestingly, the change in anisotropy due to the binding of BLT to dmDcr2^RIII^•ATP (Dr, 0.06) is ∼3-fold lower as compared to 3’ovr (Dr, 0.19; Figure 6K), suggesting a markedly reduced overall tumbling of enzyme-bound 52mer BLT dsRNA while poised for cleavage, consistent with the enhanced processivity of dmDcr-2 on BLT, compared to 3’ovr, dsRNA (Welker et al., 2011).

## DISCUSSION

A common hypothesis is that when vertebrates evolved the interferon pathway as their primary means of antiviral defense, their Dicer enzymes lost the ATP-dependent functions of their helicase domains, and became specialized for miRNA processing using their Platform and PAZ domains (Zhang et al., 2002; Fukunaga et al., 2014; Li et al., 2017). By contrast, invertebrate Dicers are key to antiviral defense, with an ATP-dependent helicase domain that orchestrates a complex reaction involving recognition of dsRNA termini, unwinding, rewinding, translocation and cleavage. While it is clear that invertebrate Dicers require ATP (Welker et al., 2011; Cenik et al., 2011; Sinha et al., 2015), how ATP coordinates Dicer activities has not been studied. Using a real time, stopped-flow method, and time-resolved fluorescence spectroscopy, we identified reaction intermediates and provided a kinetic framework for this complex molecular motor.

As illustrated, our study provides evidence of at least five intermediates for the dmDcr-2 reaction (Figure 7), and importantly, provides the first information on how these intermediates appear and decay over time. While prior equilibrium studies indicate that 3’ovr dsRNA, but not BLT dsRNA, interacts with dmDcr-2 in the absence of nucleotide, our transient kinetic studies show that dsRNA with either terminus binds to dmDcr-2 in the absence of nucleotide (Figures 1 6, and 7). However, addition of nucleotide promotes a conformational change and dynamic fluctuations that direct both BLT and 3’ovr dsRNA to the helicase domain (Figures 1, 2 and S2). Prior limited proteolysis studies (Sinha et al., 2015), as well as a cryo-EM structure with ATPgS (Sinha et al., 2015; Sinha et al., 2018), suggest that ATP binding, but not hydrolysis, is important for clamping of the helicase domain around dsRNA. Our kinetic studies are consistent with these findings, and importantly, revealed the kinetic and thermodynamic basis of termini-dependent discrimination of dsRNA clamped by the helicase domain (Figures 1 and 2). The cryo-EM structure and supporting biochemistry (Sinha et al., 2015; Sinha et al., 2018) show that dsRNA is unwound by the helicase domain. Our studies reveal the transient kinetics of this unwinding, and further, establish that the energy of ATP hydrolysis allows translocation on the unwound RNA, in a 5’to 3’ direction, prior to a slow rewinding (Figures 3 and 4). Translocation also facilitates interaction of the termini with the Platform•PAZ domain. Intriguingly, our studies suggest that ATP binding and hydrolysis by the helicase domain promote termini-dependent conformational transitions, on the nanosecond and second time-scales, ∼80Å away in the Platform•PAZ domain, and further, that these dynamics fine-tune substrate cleavage and siRNA release (Figures 5 and 6). Below we describe the salient mechanistic features of the dmDcr-2-catalyzed reaction uncovered by our studies and their implication for antiviral RNAi.

**Figure 7:**
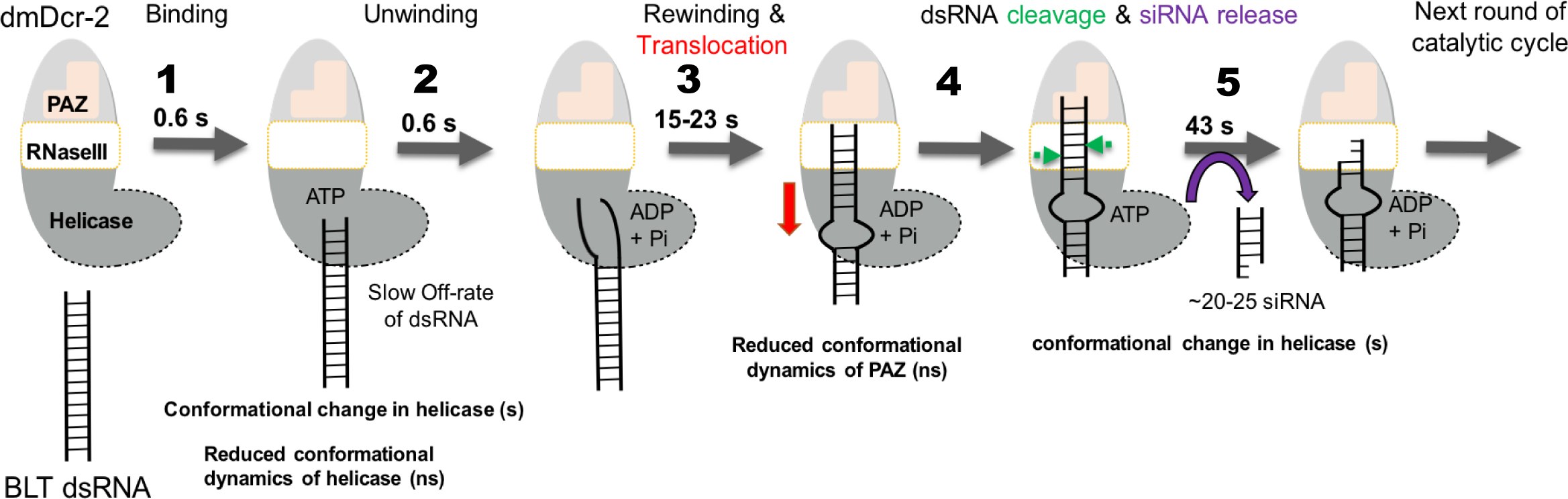
Kinetic model for ATP-dependent dmDcr-2 catalyzed reaction with BLT dsRNA. (1) ATP binding to the helicase domain promotes a closed conformational state which traps dsRNA. (2) ATP hydrolysis catalyzes dsRNA unwinding at dsRNA terminus. (3) Unwinding is followed by slow rewinding in concert with an ATP-dependent directional translocation (Red arrow) of dmDcr-2 to the Platform•PAZ domain, which directs cleavage. Platform•PAZ domains are shown in pink with a shape indicating optimal binding to 3’ovr termini compared to BLT termini. (4) The RNase III domain of dmDcr-2 cleaves (green arrowheads) dsRNA by measuring ∼21-22 nts from the terminus bound to the Platform•PAZ domain. (5) The siRNA is released via an ATP-binding mediated conformational change in the helicase domain which shows long range communication with the Platform•PAZ domain bound to siRNA termini.

### Termini-dependent dsRNA discrimination by dmDcr-2

While it is clear that *D. melanogaster* requires the helicase domain of dmDcr-2 to mount an effective antiviral response (Lee et al., 2004; Marques et al., 2013; Kandasamy et al., 2017; Donelick et al., 2020), definitive evidence that dmDcr-2’s helicase domain discriminates termini of viral dsRNA awaits in vivo validation. However, our studies reveal dramatic differences in the interaction and processing of BLT and 3’ovr termini, firmly establishing that the enzyme has indeed evolved the ability to discriminate these termini. We find that in the presence of ATP, both substrates interact with the helicase domain, and while their reaction mechanism is qualitatively similar, the kinetic parameters for their processing are quite different. The half-life for the initial encounter of BLT dsRNA with the helicase domain is an order of magnitude faster than for 3’ovr dsRNA (Figure 1I), and the rate limiting step for the observed dissociation of the enzyme-bound BLT dsRNA is 4-fold slower than that of 3’ovr dsRNA (Figure 2H). Most importantly, BLT dsRNA is processed 2-fold faster than 3’ovr dsRNA. Taken together, our studies support the idea that dmDcr-2 has evolved the ability to discriminate termini, and that this discrimination is observed at multiple steps of the catalytic cycle. This discrimination offers a mechanism whereby invertebrate Dicers could efficiently engage and process the BLT dsRNA produced by some viruses (Mueller et al., 2010; Schlee 2013; Sowa et al., 2020).

### The role of ATP binding and hydrolysis in dmDcr-2 catalysis

Our transient kinetic studies show that ATP binding promotes a termini-dependent initial encounter of dsRNA with dmDcr-2, followed by isomerization of the encounter-complex, characterized by a slow conformational change in the helicase domain. Using the *t21/2* values (Figure 1I), we calculated the observed rate constants (*kobs* =0.693/*t1/2*) for the nucleotide-dependent conformational change in dmDcr-2’s helicase domain as 0.012s^-1^ and 0.007s^-1^ when bound to BLT and 3’ovr dsRNA, respectively. Such a conformational transition would be advantageous for dmDcr-2 in trapping dsRNA during the initiation of the catalytic cycle. Our studies reveal that a differential engagement of dsRNA termini by the helicase domain bound to nucleotide offers a kinetic control mechanism for substrate discrimination. Understanding the structural basis of the nucleotide-mediated discrimination of dsRNA termini by the helicase domain awaits high resolution structures of enzyme-substrate complexes.

We find that ATP-hydrolysis by the helicase domain is essential for energetically costly steps of the catalytic cycle, including unwinding/rewinding and translocation, and conformational dynamics of the Platform•PAZ domain prior to cleavage (Fig 7). Interestingly, all of these kinetic events are 2-fold different for BLT, compared to 3’ovr, dsRNA (Figures 1, 3, 4, and 5), which is well correlated with the 2-fold difference in the rate of ATP-hydrolysis by the enzyme while bound to these substrates (Sinha et al., 2015). In addition, the time-scale for the nucleotide-mediated conformational change in the helicase domain for both BLT and 3’ovr dsRNA correlates well with siRNA release (compare *t21/2*, ATPgS, Figure 1I with *t21/2*, ATP, Figure 5E), raising the possibility of a direct involvement of the helicase domain in siRNA release. Such a role for ATP-binding during siRNA release has been proposed for human Dicer (Zhang et al., 2002). Notably, the regulatory factors Loqs-PD and R2D2 bind the helicase domain of dmDcr-2 to modulate dsRNA processing and transfer of siRNA to Argonaute-2, respectively (Liu et al., 2003; Liu et al., 2006; Marques et al., 2010; Hartig et al. 2011). We propose that an ATP-mediated conformation change in the helicase domain, in conjunction with R2D2, kinetically fine-tunes the transfer of siRNA from dmDcr-2 to Argonaute-2 at the end of the catalytic cycle.

### Is siRNA release the rate-limiting step of dmDcr-2 catalysis?

The slowest step of an enzyme-catalyzed reaction defines the rate limiting step, which is often key for regulating biological processes (Hammes and Wu, 1971). Since the half-lives for ATP-dependent cleavage of BLT and 3’ovr dsRNA, and siRNA release, are the longest steps of the catalytic cycle (Figure 7), we can unambiguously assign this as the rate limiting step. We note that the FRET-based assay does not quantitively measure the individual microscopic rate constants associated with cleavage of dsRNA and siRNA release. However, the chemical transformation, in this case cleavage of two phosphodiester bonds, is seldom the rate limiting-step in enzyme catalysis (Cleland, 1974), and it seems more likely that product release, often associated with a specific conformational change that is the reverse of substrate binding, serves as the limiting step (Cleland, 1974; Gutfreund, 1995; Fersht, 1999). Indeed, the observed rate constant associated with cleavage/siRNA release with ATP (*t21/2*, 43.2s, BLT; 83.8s, 3’ovr, Figure 5E) measured in our FRET-based assay is essentially the same as the rate of isomerization of dmDcr-2•ATPgS•dsRNA (*t21/2*, 55.5s, BLT; 95.9s, 3’ovr, Figure 1I), raising the exciting possibility that the ATP-binding-mediated conformational change in the helicase domain is critical for siRNA release. Since ATP-dependent unwinding and translocation –which are downstream of the binding event—are faster than the initial encounter of dsRNA termini with dmDcr-2• ATPgS, we propose that the free energy of ATP-hydrolysis enhances the isomerization step to funnel the isomerized intermediate to the next step of the catalytic cycle.

### The importance of conformational fluctuations of dmDcr-2 in catalysis

Conformational dynamics of distinct regions of proteins are often harnessed by enzymes to enhance catalytic efficiency (Vendruscolo and Dobson 2006; Boehr et al., 2006). Slow transitions in the micro-millisecond range are typically associated with domain movements that facilitate substrate binding and product release (Gerstein et al., 1994). In contrast, fast, pico-nanosecond conformational fluctuations promote rapid sampling of conformations that often reduce activation barriers and fine tune transient mechanisms that promote enzyme catalysis (Henzler-Wildman and Kern., 2007; Benkovic and Hammes-Schiffer., 2003). Indeed, the “jigglings and wigglings” of atoms/molecules initially envisioned by Richard Feynman seems to fine tune the transient kinetic mechanism of enzyme catalysis (Feynman et al., 1963; Benner 1989).

Our mechanistic investigation of the dmDcr-2-catalyzed reaction revealed conformational fluctuations of the helicase and Platform•PAZ domains in the nanosecond time-scale (Figure 6K) that are coupled with the free energy of ATP binding/hydrolysis by the helicase domain. Excitingly, this suggests that ATP-hydrolysis promotes a long-range communication between the helicase and Platform•PAZ domains, which are estimated to be ∼80Å apart. Our studies show that these fluctuations are termini-dependent (Figure 6), indicating these conformational dynamics are important to substrate discrimination. A direct correlation between rates of ATP-hydrolysis coupled with the reduction in conformational fluctuation of the Platform•PAZ domain (Sinha et al., 2015; Figure 6), and the ∼2-fold difference in the rate of ATP-dependent cleavage of dsRNA with different termini, further underscore the significance of a functionally-relevant fluctuation in poising the enzyme for catalysis (Figure 5). Functionally relevant and fast (nanosecond) conformational fluctuations are often coupled with longer conformational transitions (millisecond-second) in enzymes, and this often plays a direct role in substrate binding and product release (Henzler-Wildman and Kern., 2007; Gerstein et al., 1994). In this view, we propose that a markedly slow fluctuation of the helicase and Platform•PAZ domains of dmDcr-2 with BLT dsRNA, in comparison to 3’ovr dsRNA, are correlated with faster termini-dependent dsRNA binding and siRNA release. Likewise, such fluctuations are also expected to modulate the dissociation off-rate of the enzyme-bound substrate (Figure 2), and thereby impact processivity of dmDcr-2. Future studies will be needed to uncover other functionally relevant conformational fluctuations of dmDcr-2 ranging from picosecond to sec time-scales that are linked with different steps of the catalytic cycle.

## Acknowledgements

We thank members of the Bass lab for helpful discussions. This work was supported by funds to B.L.B. from the National Institute of General Medical Sciences (R01GM121706) and startup funds to R.N. from the University of Utah Chemistry Department.

## Author Contributions

RKS and BLB conceptualized the study. Transient kinetic experiments were performed by RKS. MJ and RKS performed time-resolved fluorescence experiments and RKS, MJ and RN interpreted and analyzed these data. EL provided technical assistance in RNA preparation and dmDcr-2 purification. NAR provided technical assistance in stopped-flow experiments. RKS and BLB analyzed data and wrote the manuscript with input from all authors.

## Competing Financial Interests

None.

## METHODS

### Expression and purification of dmDcr-2 and variants

dmDcr-2, dmDcr-2^RIII^, and dmDcr-2^DHel^ were expressed and purified using a baculovirus expression system as described (Sinha and Bass 2017). Briefly, recombinant pFastBac plasmid containing the open reading frame for OSF-tagged dmDcr-2 was transformed into DH10Bac competent cells. Recombinant bacmid was isolated and transfected into SF9 cells to make viral stocks (P0, P1 and P2) for protein expression. Recombinant P2 viral stock were titered and used for large-scale expression. Expressed protein was purified to homogeneity using Strep-Tactin affinity chromatography as described, and purified protein was dialyzed, concentrated, and stored at -80°C in cleavage assay buffer (25 mM TRIS pH 8.0, 100 mM KCl, 10 mM MgCl2.6H2O, 1 mM TCEP), supplemented with 20% glycerol.

### RNA preparation

BLT 52-dsRNA was prepared from 52 nt sense and antisense strands, and 3’ovr dsRNA was prepared with 52 nt sense and 54 nt antisense strands (sequences below); all strands were chemically synthesized using reagents from Glen Research (Sterling VA) (Sinha et al., 2018), and HPLC purified at the DNA/Peptide Synthesis Core facility at the University of Utah. All dsRNAs were prepared with two deoxynucleotides at the 3’end of the sense strand, and a biotin at the 5’-end of the antisense strand, to facilitate directional binding of dmDcr-2. Single strands were gel purified after PAGE (17% denaturing). dsRNA was prepared by mixing equimolar amounts of sense and antisense RNAs in annealing buffer (50 mM Tris pH 8.0, 20mM KCl), heating at 95°C for 2 min, and allowing to cool to room temperature for 4 hrs. Annealed dsRNA was purified after 8% native PAGE.

#### 52 nt sense strand for BLT and 3’ovr dsRNA

5’GGAGGUAGUAGGUUGUAUAGUAGUAAGACCAGAC CCUAGACCAAUUCAUG**<u>CC</u>**-3’

**<u>CC</u>** = deoxynucleotide

#### 52 nt antisense strand for BLT dsRNA

Biotin-5’-GGCAUGAAUUGGUCUAGGGUCUGGUCUUACUACUAUACAACCUACUACCUCC-3’

#### 54 nt antisense strand for 3’ovr dsRNA

Biotin-5-GGCAUGAAUUGGUCUAGGGUCUGGUCUUACUACUAUACAACCUACUACCUCCCC-3’

### Labeling of RNA

Cy3-5’-end-labeled 52 nt sense RNA (sequence above) was purchased from IDT (Integrated DNA Technologies). For internal labeling of sense and antisense strands, oligonucleotides were synthesized in-house with C6-amine modified uridine at desired locations needed for monitoring duplex unwinding, translocation and cleavage. The Sulfo-NHS-ester forms of Cyanine dyes (Cy3 or Cy5) were purchased from Lumiprobe (Lumiprobe corporation). C6-amine modified RNA was mixed with 20 molar excess of the NHS-ester modified dye in freshly prepared labeling buffer (100mM Sodium tetraborate pH 8.5; pH adjusted with 12.1M HCl) as described previously (Joo and Ha 2012). The mixture was incubated for 6 hours at room temperature followed by overnight at 4°C. Labeled RNA was ethanol precipitated, rinsed with 70% cold ethanol to remove excess dye, and gel purified after 17% denaturing PAGE prior to annealing to form duplex.

### Transient kinetic experiments

Kinetic events prior to dsRNA cleavage (binding, unwinding/rewinding and translocation) were investigated separately utilizing dsRNA with fluorophores (Cy3 or Cy3-Cy5) at specific sites, and by monitoring time-dependent changes in fluorophore signal. Transient kinetic analysis of dsRNA binding to dmDcr-2 was performed using a stopped-flow system (SFM 3000, BioLogic Sciences Instruments) under pseudo-first order conditions in cleavage assay buffer (25 mM TRIS pH 8.0, 100 mM KCl, 10 mM MgCl2.6H2O, 1 mM TCEP) at 25°C. The premixing concentrations of dmDcr-2 (or dmDcr-2•ATP/ATPgS) and Cy3-end labeled dsRNA in stopped-flow syringes were 2µM and 0.2µM, respectively. The nucleotide-bound form of dmDcr-2 (dmDcr-2•ATP/ATPgS) was prepared by incubating 2µM dmDcr-2 for 5 min with a large excess (8mM) of ATP/ATPgS to ensure rapid binding. The binding reaction was started with rapid mixing of equal volumes of enzyme (or enzyme•nucleotide) with dsRNA. Reaction progress was monitored by exciting the sample at 530 nm and measuring fluorescence emission using a 550 nm long pass-filter (NewPort). At least 5-10 kinetic traces were obtained for each experimental condition and values reported as averages. Averaged traces were analyzed using single or double exponential rate equations, yielding values for the observed rate constant (*kobs* = 0.693/t1/2). Data analysis was performed using the Origin software package (OriginLab corporation).

### Measurement of dissociation off-rates

Dissociation off-rates for enzyme-bound dsRNA were measured by mixing 10-fold excess of 52-dsRNA with dmDcr-2•ATPgS•Cy3-dsRNA in stopped-flow syringes. Off-rate measurements were performed in cleavage assay buffer (25 mM TRIS pH 8.0, 100 mM KCl, 10 mM MgCl2.6H2O, 1 mM TCEP) at 25°C. Dissociation kinetics were recorded by illuminating samples at 530 nm and detecting fluorescence emission using a 550 nm long pass-filter (NewPort). At least 5-10 traces were obtained and values averaged. Averaged traces were analyzed using single or double exponential rate equations to yield dissociation off-rates. Data analysis was with the Origin software package (OriginLab corporation). The activation free energy was calculated using Eyring equation (Eq. 1).

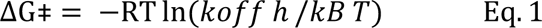

where R is the gas constant (1.986 cal K^−1^ mol^−1^), T is the absolute temperature, h is Planck’s constant (1.58 × 10^−34^ cal.s), and *kB* is Boltzmann’s constant (3.3 × 10^-24^ cal K^−1^).

### Measurement of translocation kinetics

Translocation assays used 52-dsRNA with a Cy3 covalently attached to the 18^th^ nt of the sense strand and were performed in cleavage assay buffer (25 mM TRIS pH 8.0, 100 mM KCl, 10 mM MgCl2.6H2O, 1 mM TCEP) at 25°C. Reactions were monitored upon mixing 10-fold excess of dsRNA with dmDcr-2 alone, or bound to nucleotide (dmDcr-2•ATP/ATPgS), in stopped-flow syringes. dmDcr-2•ATP/ATPgS was prepared by incubating 2µM of dmDcr-2 with 8mM ATP/ ATPgS for 5 min as described above. The time-dependent arrival of dmDcr-2 near to the Cy3 attached to dsRNA leads to PIFE (Protein Induced Enhanced Fluorescence). At least 5-10 traces were obtained and values averaged. Averaged traces were analyzed using single or double exponential rate equations, yielding values for the observed rate constant (*kobs* = 0.693/t1/2) for the translocation. Data analysis was performed using the Origin software package (OriginLab corporation).

### Measurement of dsRNA cleavage kinetics

Kinetics of dsRNA cleavage and siRNA release was investigated by monitoring the first round of catalytic cycle for ∼5 min. The reaction-mixture was synchronized at the beginning of the catalytic cycle via rapid mixing using the stopped-flow system, and by using 10-fold excess of dmDcr-2 over dsRNA. The excess enzyme over substrate ensured that dsRNA remained bound during the first turnover, thus making precise measurement of kinetic parameters feasible (Fersht 1999). dsRNA used in the cleavage assay had a FRET pair at the primary cleavage site, and cleavage and siRNA release were recorded by exciting at 530 nm (Cy3 excitation) and monitoring the time-dependent change in FRET signal (Cy5 fluorescence emission) using 670 nm long pass filter. Assays were performed in cleavage assay buffer (25 mM TRIS pH 8.0, 100 mM KCl, 10 mM MgCl2.6H2O, 1 mM TCEP) at 25°C in the absence and presence of nucleotide (ATP/ATPgS). Nucleotide-bound dmDcr-2 (dmDcr-2•ATP/ATPgS) was prepared by incubating 2µM of dmDcr-2 with 8mM ATP/ ATPgS for 5 min. At least 5-10 traces were obtained and values averaged. Averaged traces were analyzed using single or double exponential rate equations, and data analysis was performed using the Origin software package (OriginLab corporation).

### Time-resolved fluorescence experiments

#### Experimental setup

**IRF values:** The first trial was 40ps, second trial 60ps, so on average 50ps. **Beam polarization:** Vertically polarized to interrogate the sample. **Resolution:** 16ps

**Integration time:** 110s

**Sample description:** Condensed phase samples were prepared right before tested and placed into a 1mm path length cuvette.

**Laser power:** 170-175 µW

**Stability:** Stability of the beam varied by 400uW in the first trial and 1uW in the second trial.

**Wavelength and FWHM:** 528.5 nm with a FWHM of 3.5 nm.

**Wavelength ranges for filters and dichroic:** 562 short pass dichroic, 20nm bandpass centered at 586nm.

**Channel equalization with neutral density:** A 1.0 ND Thorlabs filter was added to the vertical channel’s collection fiber path to equalize the raw count rate in both polarization channels. Since both data streams are collected simultaneously, this allows for longer integration times that optimize the signal to noise ratio in the overall experiment.

#### Measurement of fluorescence lifetimes and time-resolved fluorescence anisotropy

Time-resolved photoluminescence experiments on fluorescently-labeled RNAs were used to measure fluorescence lifetime and transient fluorescence anisotropy of the Cy3 chromophore. Data were collected with a polarization-resolved epifluorescence setup whose source is the second harmonic of a tunable-wavelength pulsed Ti:sapphire laser (Coherent Chameleon Ultra II, repetition rate of 80 MHz, <200 fs pulse duration). Excitation pulse energy was kept low (2 pJ/pulse) to prevent sample photodamage, and excitation wavelength was set to 529 nm (FWHM ≤ 3.5 nm). Excitation beam polarization was set to the vertical axis with high extinction ratio polarizers before being directed to the sample with a 562 nm long-pass dichroic, and focused using a 75 mm focal length aspheric lens. Samples were held in a 4 mm pathlength quartz cuvette. Fluorescence was collected with the same lens and filtered with the same dichroic mirror before passing through a bandpass filter (576-596 nm transmission window) to further eliminate any scattered light from the excitation beam. The fluorescence signal was split into vertical and horizontal polarization channels using a set of polarizing beam splitters, and the signal in each channel was detected with identical single-photon Si photodiodes (MPD systems), whose output was routed to a time-correlated single-photon counting (TCSPC) module (PHR 800 and Picoharp 300, PicoQuant). All spectral cleanup, focusing, collection, and detection optical elements were mounted on a 30-mm cage system to prevent deleterious stray light (<1 Hz dark count rates). To match the signal-to-noise ratio in the data of each polarization channel (which have different collection efficiencies), count rates on the TCSPC module were equalized to within a factor of 2 by placing a neutral density filter (optical density = 1.0) in the path of the vertical channel. The instrument response function for setup was ≤ 60ps and data resolution was kept at 16ps. For each sample, data collection was integrated for 110 seconds. Individual traces of intensity vs. time delay for the vertical and horizontal polarization channels were stored and exported as ASCII files for further processing.

#### Data analysis (Time-resolved fluorescence)

Raw data for vertically- and horizontally-polarized fluorescence traces were imported into Matlab for processing and analysis. The temporal axis for each channel was matched to account for different sample-to-detector path lengths (∼380ps shift in time delay axis). Counts due to background signal were estimated as average of signal before the steep rise due to arrival of excitation pulses, and subtracted from data. Quantitative match of the signal collection efficiency in the orthogonal polarization channels was achieved by finding the factor g required to overlap the long-time (> 2ns) traces recorded for the free chromophore in buffer solution (tail-matching). The resulting fluorescence lifetime (300ps), anisotropy amplitude (0.40), and rotational diffusion lifetime (440ps) of this control sample (Cy3-NHS) reproduced previously-reported values (Sanborn et al., 2007). The scaling factor g for the control sample of each experimental run was used for all samples in that run. This time-shifted, background-subtracted, and scaled data were then used to calculate total fluorescence output of the sample, *I*_*tot*_ (*t*), and the transient fluorescence anisotropy, *r*(*t*), with equations 2 and 3, respectively.

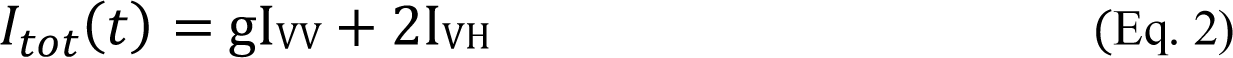

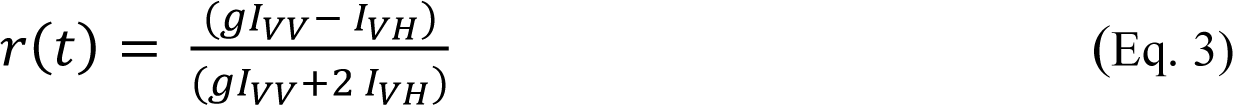

Total fluorescence output as a function of delay time was described with either a single-or bi-exponential decay using a nonlinear least-squares fitting algorithm whose output was used to calculate the sample’s average fluorescence lifetime. Transient anisotropy curves were described with an exponential decay to a constant (typically nonzero) value. Fluorescence anisotropy decay curves were analyzed using single or double exponential rate equations, and the anisotropy parameters (reported in Figure 6K) were obtained from the best of experimental data. In order to evaluate precise values of anisotropy parameters associated with sub-nanosecond dynamics, the fast phase of the decay curve was analyzed independently with a single-exponential rate equation. Goodness of fit or precision of fitted parameters was evaluated using the associated *standard error*, *Reduced Chi-square (*c*^2^*) values, as well as randomness of residuals.

### Quantification and statistical analysis

At least 5-10 kinetic traces were collected for each experimental condition used in our transient kinetic experiments. They were averaged, and the resulting averaged trace was analyzed using single or double exponential rate equations (Eqs. 4 & 5).

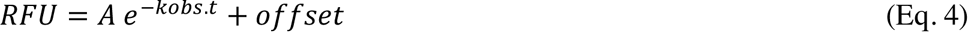

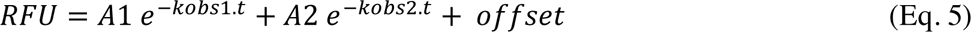

*RFU* is the relative fluorescence signal, *A* is the amplitude associated with the exponential phase, *kobs* is the observed rate constant, and *offset* is baseline signal. The relative fluorescence signal was normalized to 1 as the maximum signal for ease of comparison. The non-linear least square fitting of the data to the above rate equations were performed using Levenberg-Marquardt iteration algorithm available in Origin software package (OriginLab corporation). Goodness of fit or precision of fitted parameters was evaluated using the associated *standard error*, *Reduced Chi-square (*c*^2^*) values, and randomness of residuals.

## Supplementary Figure Legends

**Figure S1:**
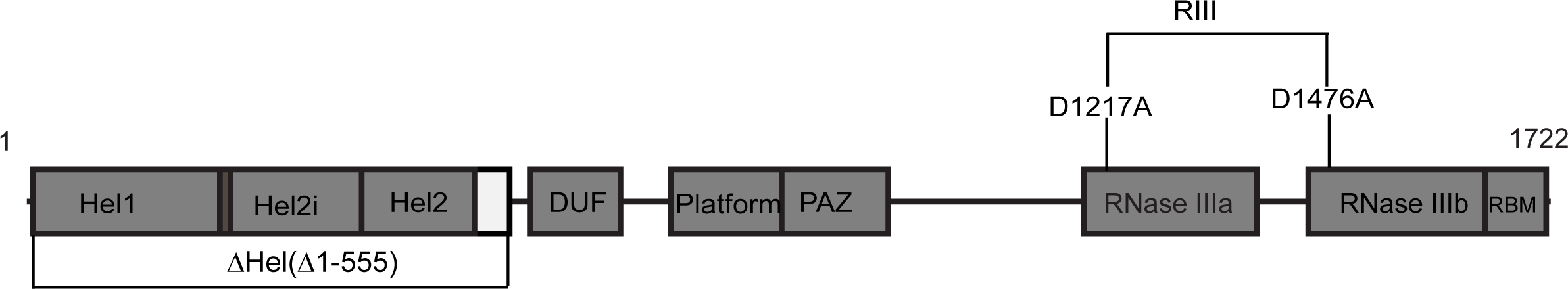
The domain organization of dmDcr-2. The boundaries of the Hel1, Hel2 and Hel2i subdomains of the helicase are indicated, as well as those for the Domain of Unknown Function (DUF), Platform, PAZ, tandem RNase III domains, and the C-terminal dsRNA binding motif (RBM). Variants used in this study are labeled.

**Figure S2.**
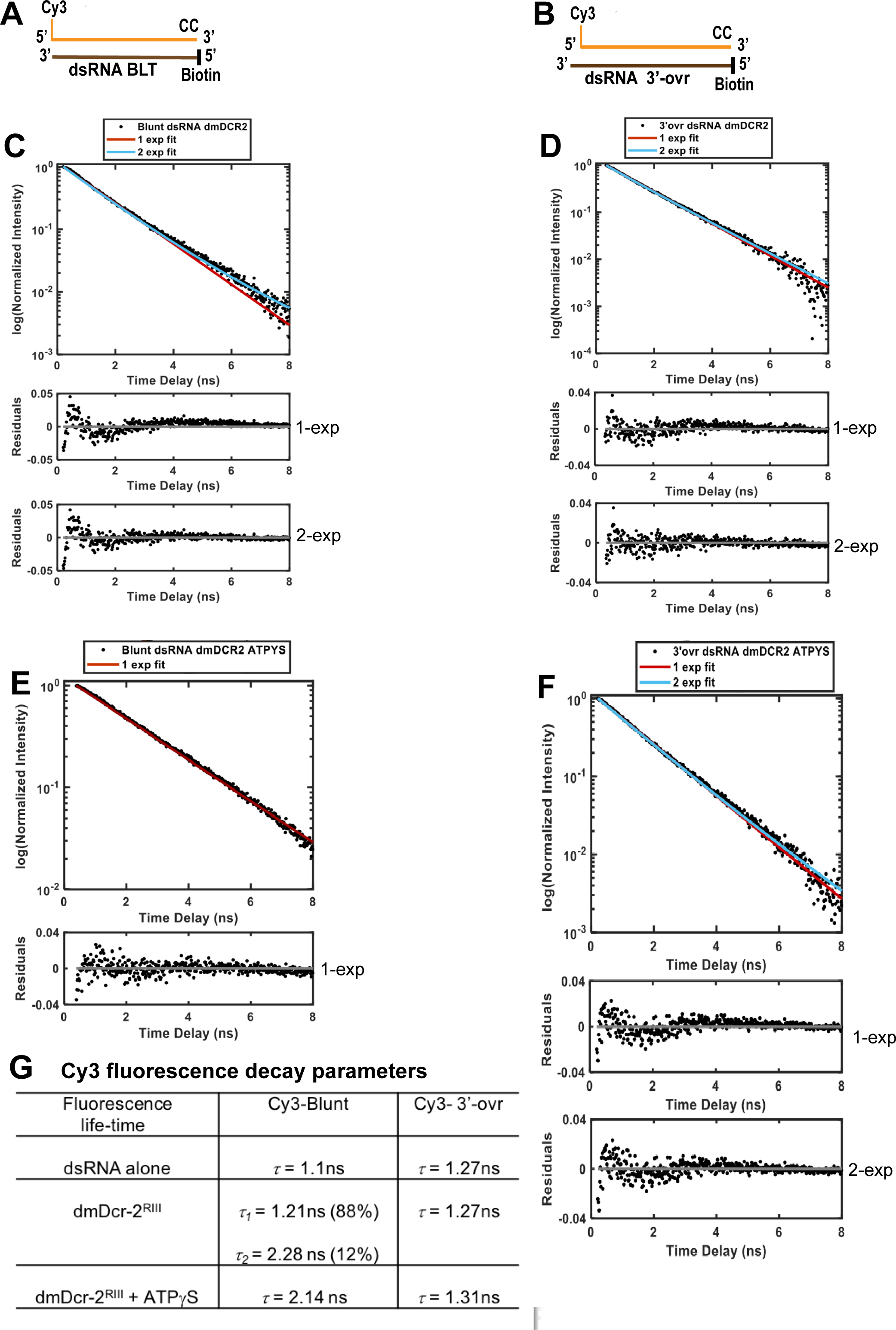
(Related to Figure 1): The fluorescence decay curve of Cy3 attached to 5’-end of BLT (**A**) and 3’ovr (**B**) 52-dsRNA while bound to dmDcr2^RIII^ (**C** and **D**) or with the dmDcr2^RIII^•ATPgS complex (**E** and **F**). dsRNA was incubated with dmDcr2^RIII^ for 5 minutes to allow complete isomerization. The RNase III variant contained two-point mutations (D1217A, D1476A), each within one of the tandem active sites that each cleave one strand (Figure S1). Data were analyzed with single and double exponential rate equations yielding values for the fluorescence lifetime (σ) and amplitude (parenthesis) of Cy3 located in different micro-environments of dmDcr-2 (**G**). Also shown are the life-time values of unbound Cy3-BLT and Cy3-3’ovr dsRNA obtained under identical experimental conditions for comparison (**G**).

**Figure S3.**
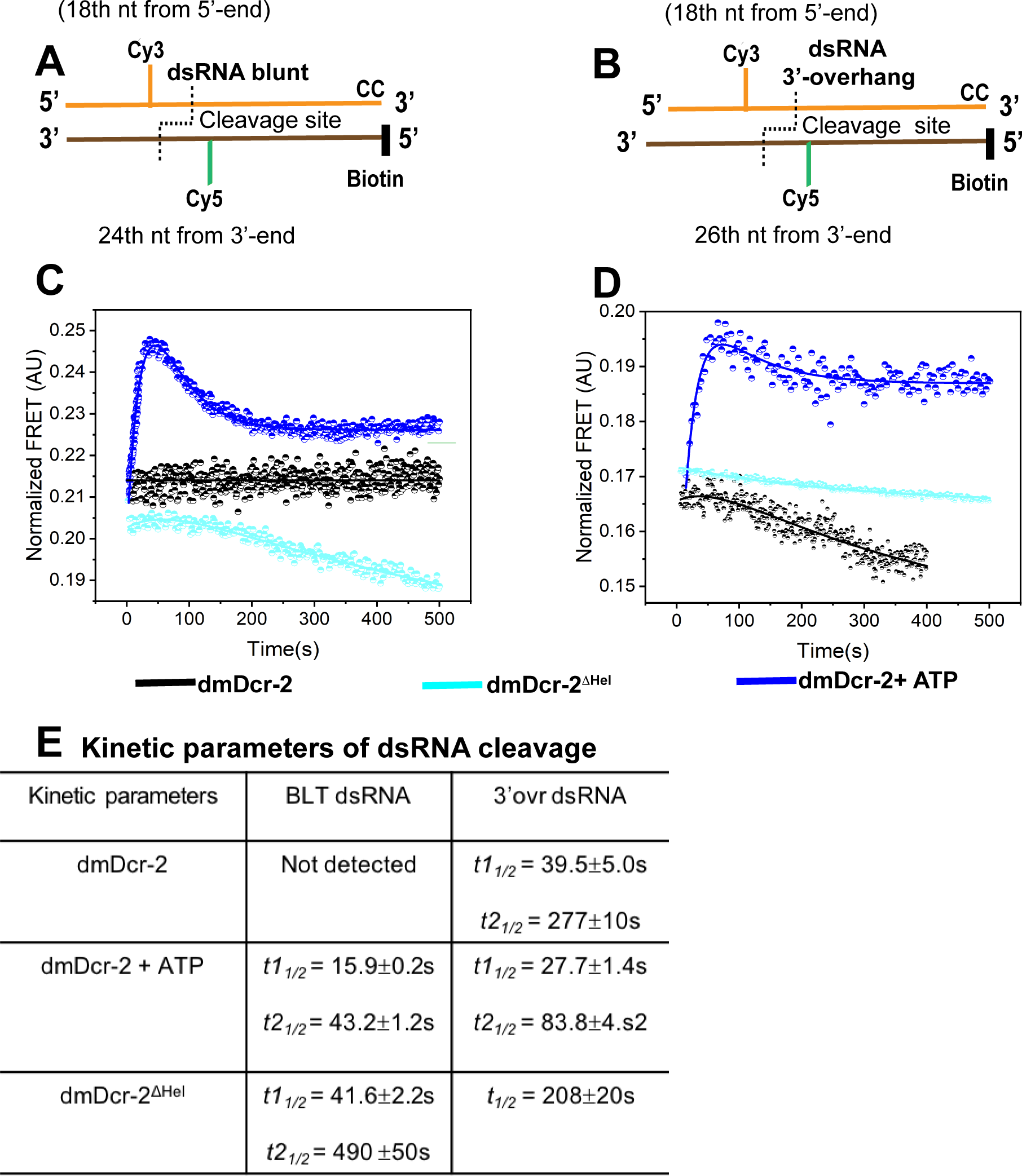
(Related to Figure 5): Stopped-flow kinetics monitoring cleavage of BLT and 3’ovr dsRNA with dmDcr2^DHel^, and for comparison, data for dmDcr-2 alone and with ATP from Figure 5 are shown. Cartoons illustrating BLT (**A**) and 3’ovr (**B**) 52-dsRNA as in Figure 5. Representative stopped-flow kinetic traces for cleavage of BLT (**C**) and 3’ovr (**D**) dsRNA by dmDcr-2 alone (black trace), dmDcr2^DHel^ (cyan trace) and dmDcr-2 with ATP (blue trace). Kinetic traces were analyzed with single or double exponential rate equations yielding half-life values (**E**).

**Figure S4.**
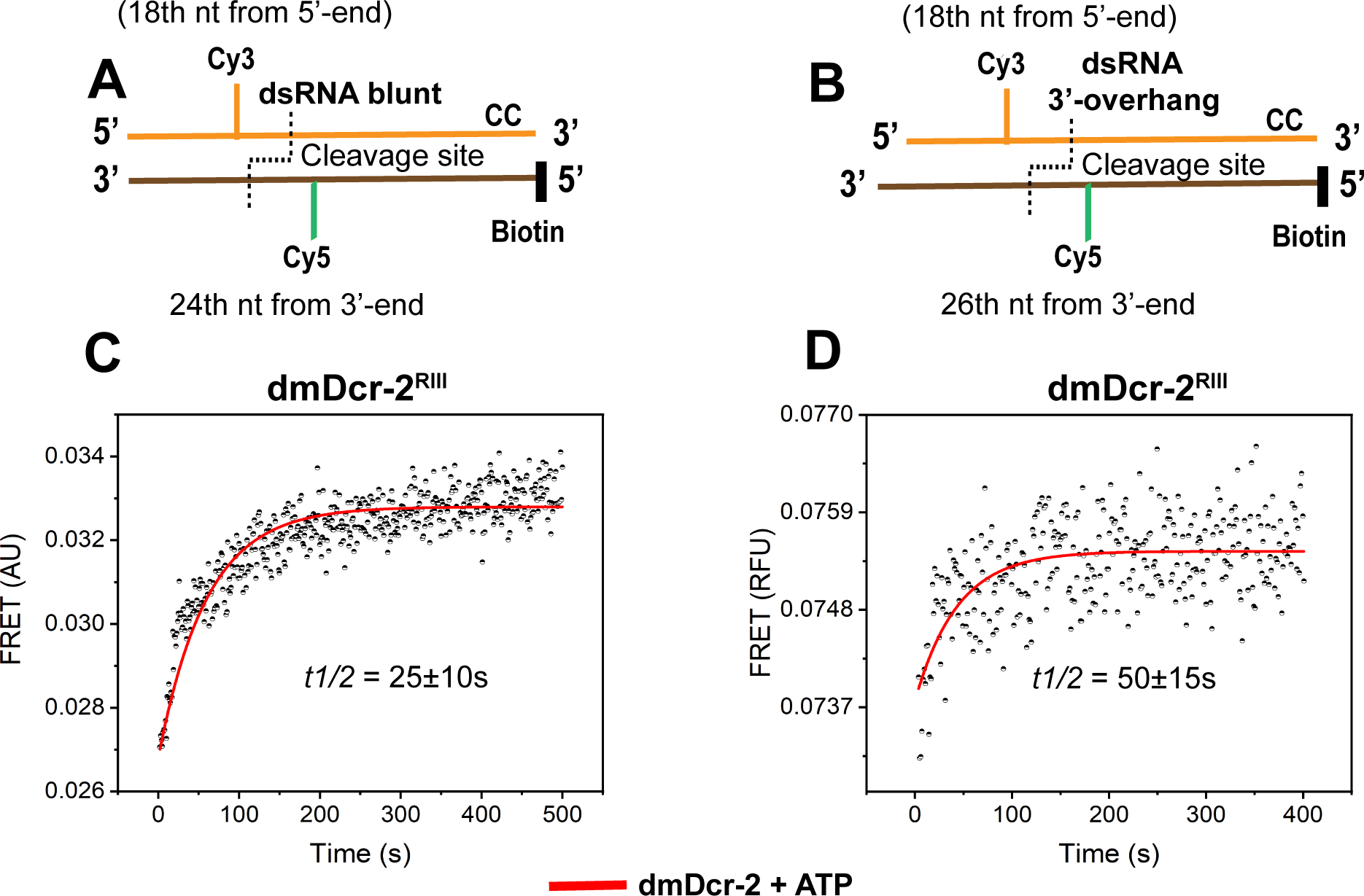
(Related to Figure 5): Stopped-flow kinetics using FRET substrates designed to monitor cleavage, but with cleavage-incompetent dmDcr-2^RIII^ in the presence of ATP. The RNase III mutant variant contains point mutations as indicated (Figure S1). Cartoons illustrate BLT (**A**) and 3’ovr (**B**) dsRNA as in Figure 5. Representative kinetic traces were analyzed with a single exponential rate equation yielding half-life values for BLT (25s) and 3’ovr (50s) dsRNA.

**Figure S5.**
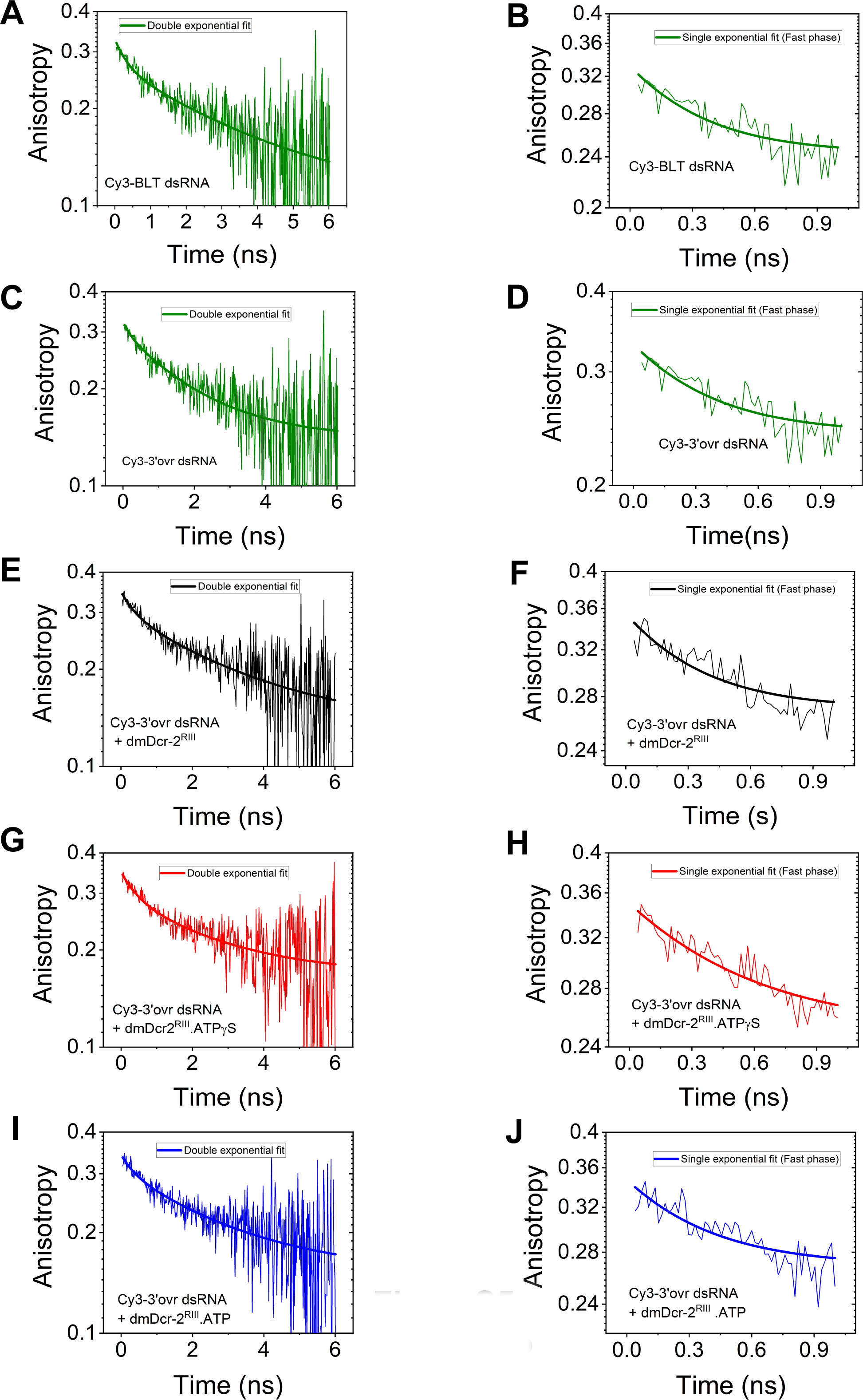
(Related to Figure 6): Fluorescence anisotropy decay curves of Cy3-labeled BLT (**A**, **B**) and 3’ovr (**C**-**J**) 52-dsRNA, -/+ enzyme and nucleotide as indicated. All biphasic anisotropy decay curves were analyzed using double-exponential equations, and to obtain precise values of anisotropy parameters associated with fast dynamics, the fast phase of decay curves was independently analyzed with a single-exponential equation, yielding anisotropy parameters listed in Figure 6K.

**Figure S6.**
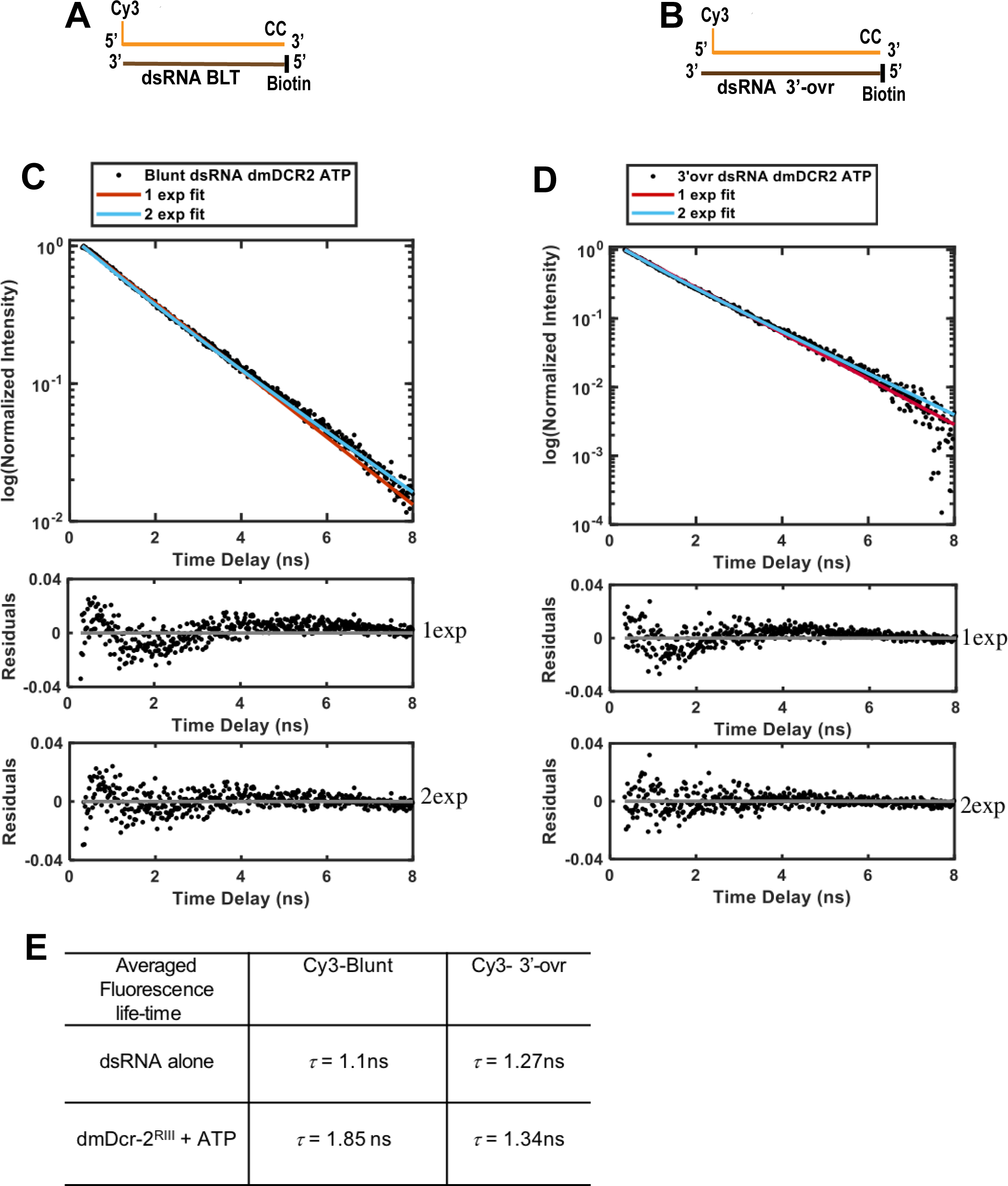
(Related to Figure 6): The fluorescence decay curve of Cy3 attached to 5’-end of BLT (**A**) and 3’ovr (**B**) 52-dsRNA while bound to dmDcr2^RIII^ •ATP (**C** and **D)**. Data were analyzed with single-and double-exponential rate equations yielding values for the fluorescence lifetime (σ) of Cy3 in different micro-environments of dmDcr-2 (E). Also shown are the life-time values of Cy3-BLT and Cy3-3’ovr dsRNA without protein (alone) obtained under identical experimental conditions for the ease of comparison (**E**). The life-time values listed here represent the averaged life-time.

## Notes

### Competing Interest Statement

The authors have declared no competing interest.

